# A polymerase ribozyme that can synthesize both itself and its complementary strand

**DOI:** 10.1101/2024.10.11.617851

**Authors:** Edoardo Gianni, Samantha L. Y. Kwok, Christopher J. K. Wan, Kevin Goeij, Bryce E. Clifton, James Attwater, Philipp Holliger

**Affiliations:** MRC Laboratory of Molecular Biology; Cambridge, CB2 0QH, United Kingdom; The Francis Crick Institute; London, NW1 1AT, United Kingdom; Department of Chemistry, UCL; London, WC1H 0AJ, United Kingdom

## Abstract

The emergence of a chemical system capable of self-replication and evolution is a critical event in the origin of life. RNA polymerase ribozymes could constitute such a system, but their large size and structural complexity hinder their self-replication and make their spontaneous emergence improbable. Here we describe QT45: a 45-nucleotide ribozyme, discovered from a random sequence pool, that catalyzes general RNA-templated RNA synthesis using trinucleotide triphosphate substrates. QT45 can synthesize both its complementary strand from a mix of all 64 trinucleotides and a copy of itself using 13 defined trinucleotides and one hexamer as substrates. The discovery of this complex activity in a small ribozyme suggests that polymerase ribozymes may be more abundant in RNA sequence space than anticipated, thereby facilitating the emergence of self-replication.

## Main Text

A hallmark of life is the replication of the genetic material with variation on which natural selection can act (*1*). How this capacity for heredity and evolution first emerged is unknown. The RNA world hypothesis posits that a catalytic RNA sequence (ribozyme) capable of driving its own replication emerged from random-sequence pools of RNA oligomers formed by prebiotic chemistry on the early Earth (*2–6*). This same heterogeneous pool of oligomers could be used by the ribozyme as substrates for self-replication (*7, 8*), starting the propagation of the genetic material within an evolving system.

To persist, such a ribozyme would need to act as an RNA-dependent RNA polymerase, catalyzing the two steps of a self-replication cycle — the synthesis of its complementary strand (-strand) and of itself (+ strand) — using general monomer or short oligomer substrates that allow the free sequence variation needed for open-ended evolution. A single RNA molecule must fulfil the multiple interdependent functions of such an RNA polymerase: general template and substrate binding, iterative phosphodiester bond formation, and accurate RNA-templated RNA synthesis. Ultimately, performing these functions with sufficient yields and fidelity to overcome chemical and mutational decay would allow it to propagate its genetic information and evolve towards higher complexity (*6, 9*).

This ability for RNA self-replication has not been observed in nature, but RNA sequences that catalyze RNA-templated RNA synthesis using mono- or trinucleotide substrates have been discovered by *in vitro* evolution and RNA engineering (*10–18*). However, while these RNA polymerase ribozymes (RPR) show how RNA can fulfil many of the functions required for self-replication, they fall short of self-replication, unable to synthesize their (+) or (−) strands even individually. This may be due to both their size (∼150-300 nt), which imposes a high synthetic burden, and their structural complexity, as stably folded RNA can pose a significant obstacle to replication (*13, 14*).

The size and complexity of these polymerase ribozymes not only reflects their descent from augmentation of the already large class I ligase ribozyme (*11, 13*) but is also in accord with the notion that functional sophistication in RNA scales with size and structural complexity (*19–21*). This raises obvious questions regarding the RNA World hypothesis, due to the implausibility of the spontaneous emergence of such large ribozymes (*22–24*). RNA sequences of the size of existing polymerase ribozymes would be both vanishingly rare in even very large sequence pools and far outside the range of RNA oligomer lengths observed to form abiotically (*25–29*). This leads to seemingly paradoxical demands, whereby ribozymes must be large and complex to encode polymerase activity, but that size and complexity impede both replication and emergence.

### Discovery of new small ribozyme motifs

To resolve these conflicting demands, we wondered if polymerase activity might be found in shorter RNA motifs. This would be advantageous both for self-replication and for emergence, as shorter sequences would be both easier to copy and more readily generated by prebiotic chemistry. To define a minimal length of RNA sequence required for polymerase ribozyme function, we carried out a *de novo* selection from a random sequence pool for the templated polymerization of RNA.

To ease the phenotypic requirements and isolate the shortest motif possible, we leveraged our past observations that RNA polymerase activity is boosted by freezing in eutectic ice, which stabilizes ribozymes and concentrates substrates (*30*). Of the range of RNA substrate lengths that may have been available prebiotically, we focused on trinucleotide triphosphate substrates (henceforth called triplets), which enable the copying of highly structured RNA templates (*13, 31*) and inhibit strand reannealing in replication cycles (*32*). Triplet substrates reduce the phenotypic requirements of a polymerase without compromising the free sequence variation in synthetic products, a prerequisite for open-ended evolution in RNA self-replication.

We initiated selections from three small, random sequence pools (∼1×10^12^ unique RNA sequences) each containing a short (20, 30 or 40 nucleotides) randomized region constituted as a tandem repeat (fig. S1). We challenged the pools first to catalyze a single templated ligation of a primer to an adjacent triphosphorylated substrate that is covalently linked to the library via a flexible RNA linker (Fig. 1A). Once catalytic activity was observed, we challenged the pool to catalyze triplet polymerization (Fig. 1B). Due to the flexible linker, active members of the pools react intramolecularly, extending the primer and ligating it with the tethered substrate. This results in a covalent coupling between active members of the library and the biotinylated primer, enabling selective recovery of active library members via streptavidin pull-down. To ensure generality of polymerase activity, selective pressure was gradually increased by requiring the templated incorporation of an increasing number of triplets and varying the template and primer sequences used throughout the rounds (table S1).

**Fig. 1.**
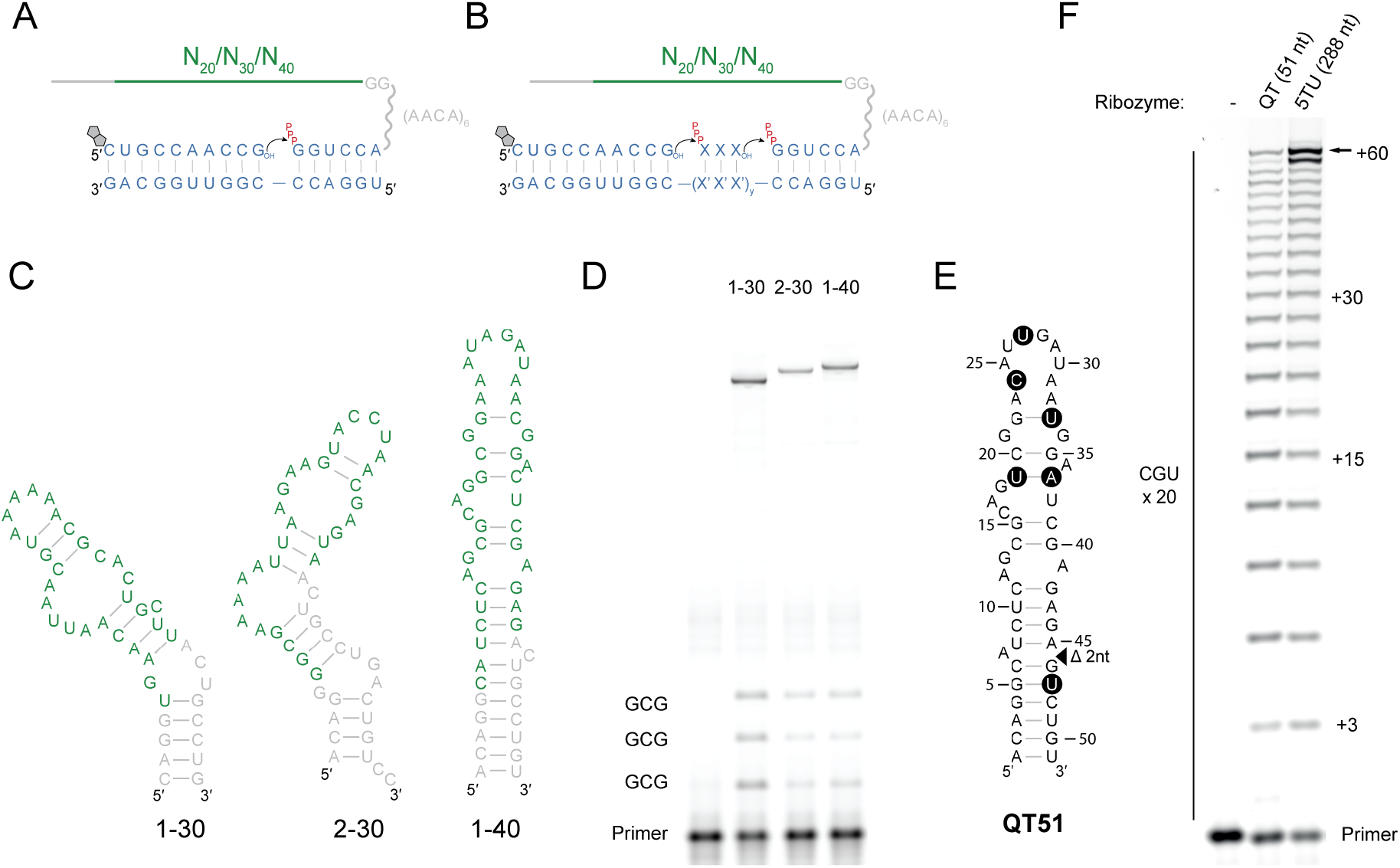
Discovery and evolution of three small polymerase ribozyme motifs. (**A**) Format of the selection construct used for the initial selection rounds (rounds 1 to 3 or 1 to 5), with the library tethered via a flexible linker to a hexameric tag that hybridizes to the template. The biotinylated primer enables the capture of active ligases (described more in detail in fig. S1-S2). (**B**) Format of the selection construct used in the later rounds (3 to 11 or 5 to 11), requiring the polymerization of triphosphorylated trinucleotide (triplet) substrates. Sequence of the triplet (XXX) and number of triplets (y) encoded by the template (X’X’X’) was varied over the course of the selection (details in table S1). (**C**) Sequence and predicted secondary structures of three ribozymes discovered from the libraries that display iterative triplet ligation i.e. triplet polymerase activity. In green, the nucleotides derived from the random library section. In grey, nucleotides derived from the constant regions (linker and primer binding site). (**D**) Iterative triplet polymerization by the ribozymes displayed in (C) in the selection construct displayed in (B), with XXX = GCG and X’X’X’ = CGC and y = 3. Reaction conditions: 50 nM ribozyme-substrate, 50 nM primer BCy3P10GA, 50 nM template t6FP10gaGCG3, 5 μM pppGCG triplet, 0.05% Tween 20, 200 mM KCl, 50 mM MgCl_2_, 50 mM CHES-KOH, pH 9, 3 days incubation at -7 °C frozen. Ribozymes are hybridized to the template. (**E**) Sequence and predicted secondary structure of the QT51 ribozyme derived from the 1-40 clone. Black circles indicate the 6 residues mutated from the 1-40 ancestral sequence; triangle indicates a 2-nucleotide deletion. (**F**) Synthesis of a 60-nucleotide sequence composed of 20 repeats of the CGU triplet. Reaction conditions: 0.25 μM primer F10, 0.25 μM template tP10CGU20, 0.25 μM ribozyme, 10 μM pppCGU triplet, QT51 in 0.05% Tween 20, 50 mM MgCl_2_, 50 mM CHES-KOH, pH 9, 5TU+t1.5 in 200 mM MgCl_2_, 50 mM Tris-Cl, pH 8.3, 2 weeks at -7 °C frozen. Ribozymes are not hybridized to the template.

We had initially designed our library construct as a tandem repeat based on the reasoning that a larger, dimeric RNA would be more likely to be able to form a complex ribozyme structure, while requiring only the monomer sequence to be replicated. However, when we assayed for polyclonal polymerase ribozyme activity in monomer and dimeric form, activity was observed in the library even in its monomer form in round 8 (fig. S2). Further rounds of selection were therefore carried out using the monomer form (fig. S3). After 11 rounds of selection, we identified three small, unrelated RNA motifs from two of the libraries (named 1-30, 2-30, 1-40) each with template-dependent RNA polymerase ribozyme activity. All three motifs satisfied the minimum requirements of iterative, cognate triplet addition for different template sequences (Fig 1D, fig. S4) with regiospecific formation of the canonical 3′-5′ phosphodiester bond (fig. S5).

### QT: a new small ribozyme class with complex RNA polymerase activity

We were surprised to find this complex activity, previously only observed in much larger ribozymes, in quite tiny RNA motifs. Each motif was subjected to mutagenesis (24% per base randomization) and 7 more rounds of selection (table S2), resulting in a dominant clone with robust triplet polymerase activity (Fig. 1E) derived from the 1-40 ribozyme. This 51-nucleotide ribozyme (named QT51) catalyzes templated phosphodiester bond formation from a 3′-OH and an adjacent 5′-triphosphate (5′-PPP) at an apparent rate (k_obs_) of 0.06 min^-1^ (fig. S6) and can copy RNA templates longer than itself, such as a 60-nt template of ACG tandem repeats (Fig. 1F). This compares favorably with 5TU+t1.5 (hereby referred to as 5TU) (*13, 31*), a previously discovered RNA polymerase ribozyme, descendant of the class I ligase, more than 5 times larger than QT51 (fig. S7), used as a benchmark of triplet polymerization throughout this study. Polymerase ribozyme activity persisted in progressively truncated (45, 40, 35 nt) versions of QT51. The 45-nt version, named QT45, retained full RNA polymerase activity (as judged by synthesis of a 42 nt product), whereas the further truncation variants QT40 and QT35 (consisting solely of a core of 29 nucleotides supported by a 3 base pair stem) retained polymerase activity but with reduced efficiency (Fig. 2A).

**Fig. 2.**
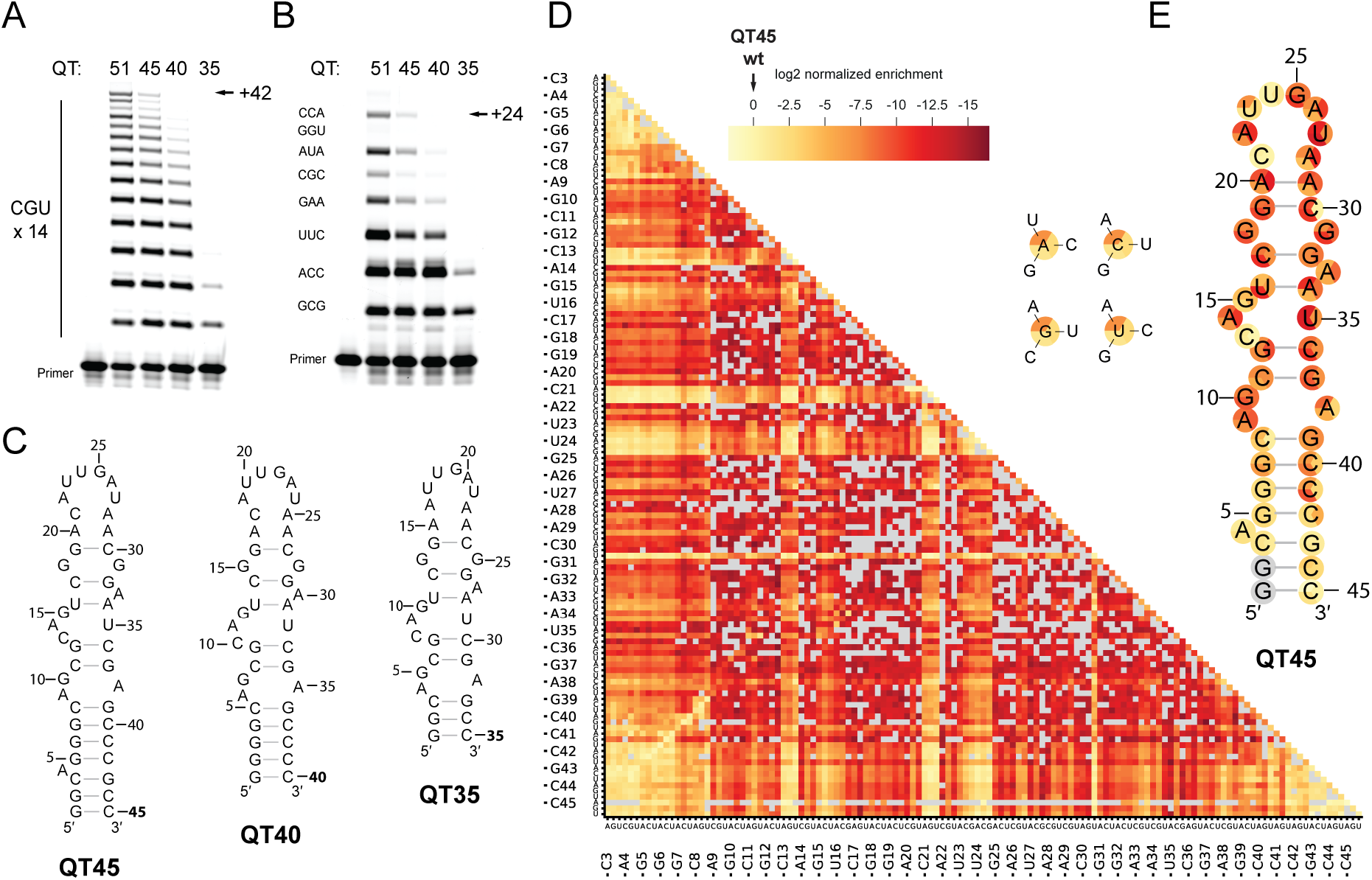
A small RNA motif with a functionally dense core encodes the triplet polymerase activity of the QT ribozyme. (**A**) Synthesis of 42 nt CGU repeat sequence by QT51 and its truncation variants. Reaction conditions: 0.25 μM primer F10, 0.25 μM template tP10CGU14, 0.25 μM ribozyme, 5 μM pppCGU triplet, 0.05% Tween 20, 50 mM MgCl_2_, 50 mM CHES-KOH, pH 9, 5 days at -7 °C frozen. (**B**) Synthesis of a mixed sequence template by the same ribozymes as in (A). Reaction conditions: 0.25 μM primer F10, 0.25 μM template t6FP10mix, 1.25 μM ribozyme, 5 μM each triplet, 0.05% Tween 20, 50 mM MgCl_2_, 50 mM CHES-KOH, pH 9, 5 days at -7 °C frozen. In both (A) and (B), ribozymes are not hybridized to template. (**C**) Predicted secondary structure diagrams of the ribozymes used in (A) and (B) (**D**) Heatmap of the primer extension activity for all measured single and double mutants of the QT45 ribozyme, with the first constituent point mutation indicated on the x-axis and the second one on the y-axis. Missing datapoints are shown in gray. (**E**) Predicted secondary structure of QT45 ribozyme with nucleotide color corresponding to its measured average activity on three different templates for each of the three possible mutations, in the same order as the 3 representative mutations provided, indicated in the same color scheme as (D).

QT ribozyme polymerase activity is not limited to trinucleotide triphosphate substrates but extends to the incorporation of dinucleotides and longer oligonucleotide triphosphates (fig. S8, fig. S9). Unlike the class I derived polymerase ribozyme 5TU, the QT ribozyme showed activity with 5′-adenylated substrates (fig. S9), a major side product (considered an inhibitor of nonenzymatic RNA copying) of imidazole-based prebiotic activation chemistries (*33, 34*). Such promiscuity in substrate length and chemistry might be beneficial to RNA replication in a heterogeneous prebiotic environment (*35*).

To test its general RNA polymerase activity, we challenged QT51 and its truncation variants to copy a 24 nt mixed sequence template comprising a representative variety of triplet junctions (8 of 16 possible ones). Both QT51 and its QT45 truncation could successively incorporate the 8 varying triplets required to synthesize this sequence (Fig. 2B). Although selected in tethered form linked to its substrate, QT ribozymes do not require any tethering or base pairing to the primer–template–substrate complex for RNA polymerase activity and show multi-turnover activity (fig. S10). This suggests that this ribozyme class can engage with the primer–template–substrate complex purely via general, sequence-independent, tertiary contacts. This advanced phenotype was previously only observed in the much larger t5+1 and 38-6 ribozymes and their derivatives (*13, 31, 36, 37*), or in a cross-chiral ribozyme polymerase (*38, 39*).

To better understand the basis for QT ribozyme activity and the nature of its tertiary primer–template-substrate interactions, we mapped the 2′-OH moieties needed for efficient catalysis on the five nucleotides upstream and downstream of a model ligation junction (−5, 0, +5) by 2′-deoxynucleotide substitutions (similarly to (*36, 40*)), in comparison to the 5TU ribozyme (fig. S11). For the template strand, suppression of ligation was observed for both QT45 and 5TU by 2′-deoxy-substitutions at positions - 4 and +2. For the primer strand, suppression of ligation was observed for 2′-deoxy-substitutions at positions -1, -2 and -3 for QT45, and positions -1 and -2 for 5TU. The overlap between tertiary contacts for QT45 and the unrelated 5TU ribozyme is unexpected but consistent with the similar phenotype of the two ribozymes and may suggest a common mode of primer-template engagement for polymerase ribozymes.

To understand the sequence determinants of polymerase function in this small RNA motif, we performed a comprehensive fitness landscape analysis on QT45, similar to a previous analysis on 5TU (*31*). We quantified the changes in genotype abundance before and after a single round of selection for triplet polymerase activity on three different templates (encoding 3 UGC, 3 AUA or 12 CUA triplets), which provided fitness estimates of all QT45 single mutants and 85% of QT45 double mutants (Fig. 2D, Fig. 2E). This revealed a sharp fitness peak, with almost all mutations being detrimental to activity. Only 1% of observed single and double substitutions were within 90% of QT45 wild-type fitness, a significantly lower fraction when compared to the fitness landscape of 5TU (*31*), where almost 9% of mutants were within 90%. Mutations in the basal stem region of QT45 were generally tolerated, especially when base pairing was maintained. In the core of QT45, only a few positions tolerated mutations. Similar trends were observed in the fitness landscape of QT39 – a variant of QT45 that maintains the same core with a truncated stem (fig. S12). This low tolerance for mutations likely reflects a high density of functional residues in the QT ribozyme core required for sustaining the multiple functional requirements of a polymerase ribozyme in a small RNA motif.

Mapping the number of functional residues present in the QT ribozyme motif allows an estimate of the likelihood of its emergence from random sequence pools. Of the 30 nucleotides present in the core motif, 3 tolerated all point mutations without substantial effects on fitness (C13, C21 and U24), 2 tolerated mutations to another residue (U23 and C30) and a 4 base pair basal stem could support polymerase activity as long as the stem was not disrupted. The intrinsic probability of finding this specific fold in random sequence is thus ∼4.4×10^−18^ (calculated as in (*19*)). The probability of identifying the QT ribozyme fold is likely an underestimate of the actual abundance of these phenotypes in sequence space, given the identification of multiple triplet polymerase ribozymes from a starting pool of only ∼1×10^12^ molecules (albeit further mutagenized over the course of the selection), suggesting that they are more abundant than anticipated and may be within reach of abiotically-available RNA pools.

### Ribozyme-catalyzed synthesis of an active ribozyme

To further examine the general polymerase activity of the QT ribozyme, we challenged QT51 to synthesize a functional RNA sequence, specifically a minimal 18 nt hammerhead endonuclease ribozyme (HHz) (*41*). QT51 was able to synthesize the HHz to full length using the six defined triphosphorylated triplet substrates making up the HHz sequence (GCU, CGA, CUG, AUG, AGG, CGC), or using a random substrate pool comprising all 64 triplets (NNN) (Fig. 3A).

**Fig. 3.**
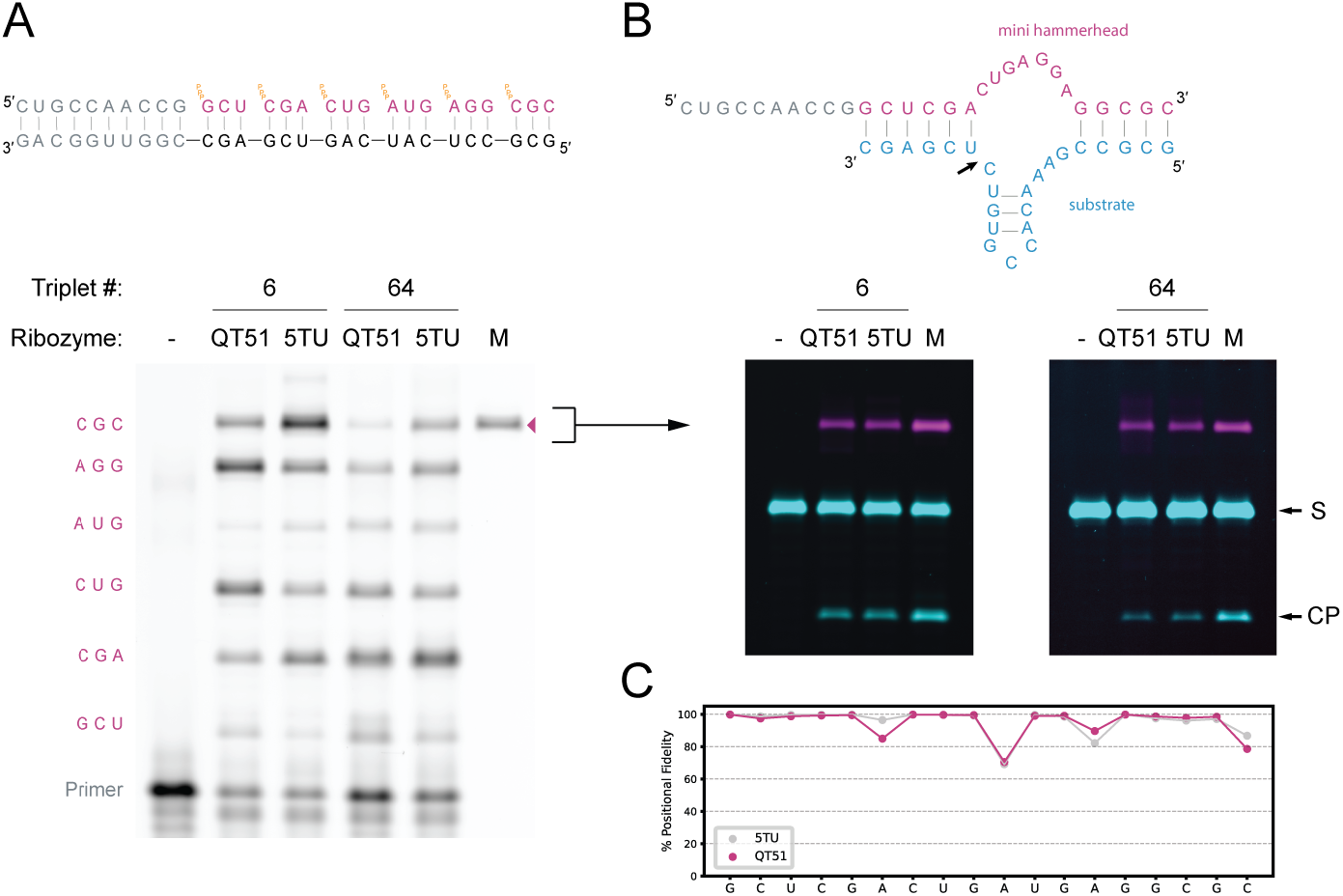
QT ribozyme-catalyzed synthesis of an active ribozyme. (**A**) (Top) Diagram of primer, template and triplets necessary for the synthesis of the mini hammerhead ribozyme. (Bottom) Ribozyme catalyzed synthesis of the mini hammerhead ribozyme. Full-length product indicated by a magenta triangle. M indicates marker full-length product lane. Reaction conditions: 0.5 μM primer BCy3P10, 0.5 μM template tP10HHz, 5 μM QT51 or 0.5 μM 5TU+t1.5, 5 μM each defined triplet or 1.75 μM each of the 64 possible triplets (NNN), 0.05% Tween 20, 50 mM MgCl_2_, 50 mM CHES-KOH, pH 9, 52 days at -7 °C frozen. (**B**) (Top) Diagram of the mini hammerhead ribozyme (magenta) in complex with its substrate (cyan). The cleavage site is indicated by an arrow. (Bottom) Cleavage activity of ribozyme synthesized hammerhead ribozymes, compared with protein polymerase synthesized controls (M lane). Synthesis reaction of the hammerhead used for cleavage was carried out as in (A) but for 36 days. Cleavage reaction conditions described in SI methods. (**C**) Positional fidelity of copying by QT51 shown in magenta, 5TU control shown in grey. Synthesis reaction was carried out as in (A) but for 28 days.

Ribozyme-catalyzed synthesis of functional RNAs (ribozyme or aptamer) is an important function in primordial RNA metabolism (*6*) and has previously only been achieved by the much larger class I ligase-derived polymerase ribozymes (*12–14*). QT51-synthesized HHz products, both from defined and from random substrate triplets, were catalytically active with cleavage activities comparable to the 5TU synthesized HHz (Fig. 3B). We analyzed QT51 fidelity by sequencing the HHz product made in the presence of all 64 triplets and compared it to the same product synthesized by the 5TU polymerase ribozyme (Fig. 3C) under the same conditions. We observed an average per nucleotide fidelity for full-length products by QT51 of 93.4%, and 94% for 5TU, with mismatches mostly due to 3 mutation hotspots shared by both ribozymes. The poor fidelity at these hotspots is due to a tolerance for a G:U wobble pair in place of the correct A:U pair, in particular at the first position of the triplet.

### Ribozyme-catalyzed synthesis of its complementary strand and of itself

The robust and general triplet polymerase activity of the QT class ribozymes, combined with their small size, suggested that they might meet the critical synthetic requirements for self-replication: the templated synthesis of both a ribozyme’s complementary strand (-strand) and of itself (+ strand) (Fig. 4A). In order to test these two reactions, we used the QT ribozyme truncated to length 45 (QT45), the smallest variant with uncompromised activity.

**Fig. 4.**
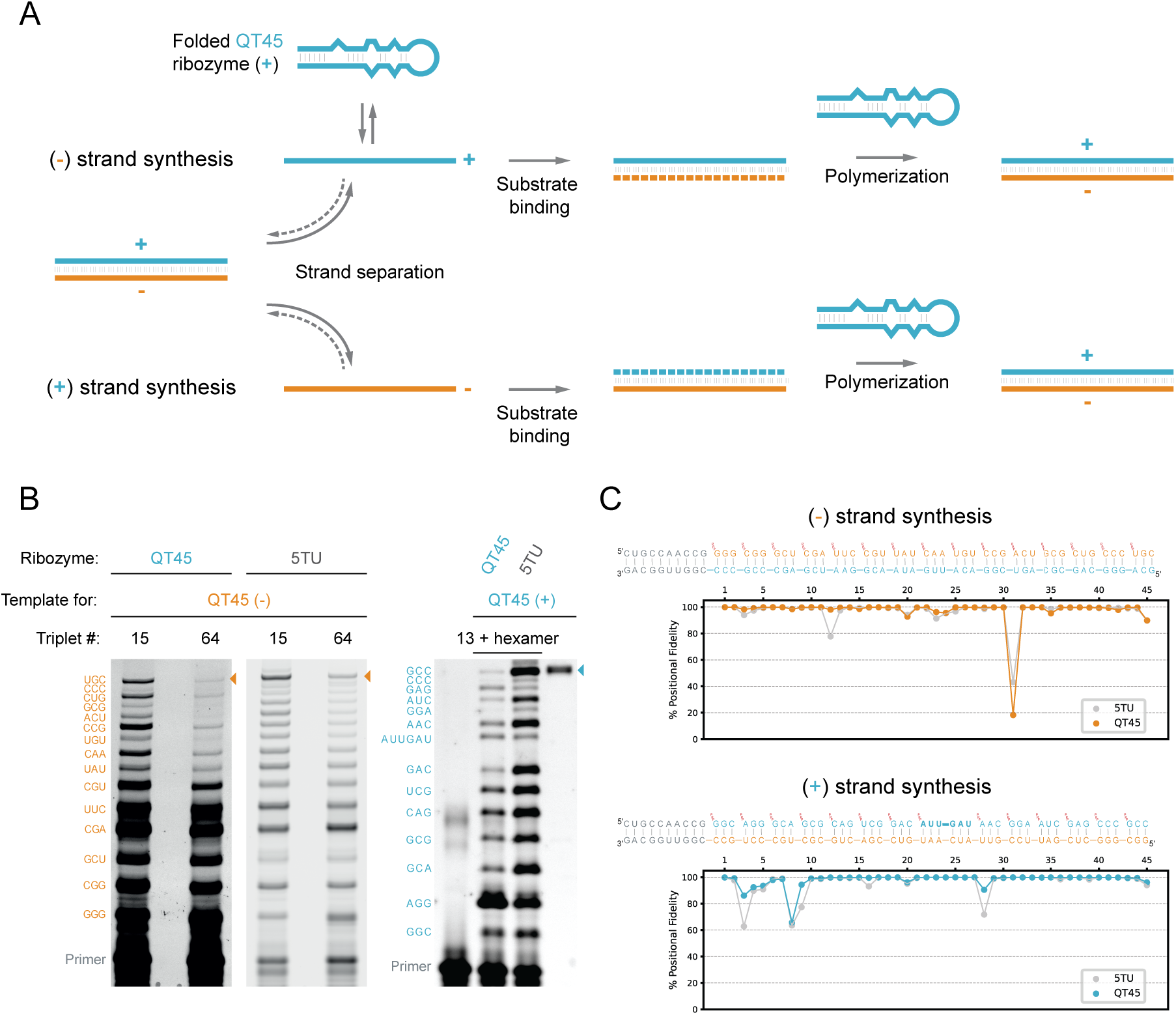
QT ribozyme-catalyzed synthesis of its complementary (-) strand and of itself (+). (**A**) Diagram of an RNA self-replication cycle: a starting duplex composed of (+) strand (corresponding to the ribozyme), and (-) strand (the reverse complement of the ribozyme), are separated. Triplets bind to the strands preventing their re-annealing. QT-catalyzed polymerization of the triplet substrates leads to the synthesis of new (-) and (+) strands. (**B**) (Left) QT45-catalyzed synthesis of its own complementary strand (QT45(-)) starting from 15 defined substrates or all 64 substrates, compared with the 5TU-catalyzed synthesis of the same sequence. Full-length product indicated by a tan triangle. Reaction conditions: 0.25 μM primer BCy3P10, 0.25 μM template t4msP10QT45, 10 μM QT45 or 0.5 μM each 5TU and t1.5, 2.5 μM each of the 15 defined triplets or 1.75 μM each of the 64 triplets, 0.05% Tween 20, 50 mM MgCl_2_, 50 mM CHES-KOH, pH 9, 28 days at -7 °C frozen. (Right) QT45 catalyzed synthesis of itself (QT45(+)) using a mix of triplet substrates with the aid of one pre-formed hexamer. Full-length product indicated by a teal triangle. Reaction conditions: 8 nM primer BCy3P10, 8 nM template t4psP10QT45, 16 nM QT45 or 5TU+t1.5, 200 nM each triplet, 0.01% Tween 20, 0.4 mM MgCl_2_, 0.6 mM KCl, 1 mM CHES-KOH, pH 9, acid-heat-cycled once, incubated for 33 days at -7 °C frozen. (**C**) (Top) Positional fidelity of copying of full-length QT45(-) from reactions in (B), with QT45-catalysed synthesis shown in tan, 5TU control shown in grey. (Bottom) Positional fidelity of copying of full-length QT45(+) from reactions in fig. S19. QT45-catalyzed synthesis shown in teal, 5TU control shown in grey.

We first explored the QT45-catalyzed synthesis of its own complementary (-) strand, wherein QT45 (the (+) strand) acts both as the template strand and as the catalyst. This requires the reconciliation of two seemingly paradoxical requirements: within the same reaction, QT45 must fold to act as a catalyst, while also unfold to act as a template. This paradox can be reconciled within the behavior of an RNA ensemble: two RNA conformations are present simultaneously within the ensemble, through a folding equilibrium. As the triplet substrates selectively bind to the unfolded template strand, this equilibrium is sensitive to the triplet concentration (fig. S13), as observed previously for templates containing secondary structures (*13*). At higher triplet concentrations, the unfolded QT45 (+) template strand conformation will be increasingly populated as triplets cooperatively bind to the (+) strand, consistent with the inhibition of QT45 by high NNN concentrations (fig. S14). At lower triplet concentrations, the folded, active QT45 (+) strand ribozyme conformation predominates.

We incubated QT45 and a QT45 with an external primer binding site in the presence of an optimal concentration of either the 15 triplets encoding its complementary (-) strand, or all 64 possible triplets (NNN). Synthesis of the full-length complementary (-) strand was observed using both defined triplets and the NNN substrate pool (Fig. 4B). Deep sequencing confirmed the identity of the full-length (-) strand synthesized using the NNN pool with an average per nucleotide fidelity of 94.1% (Fig. 4C), similar to the fidelity of HHz synthesis (Fig. 2C).

Having shown that QT45 could synthesize its own QT45 (-) template strand, we next sought to explore if it could also synthesize itself, i.e. another QT45 (+) strand from a QT45 (-) template. This again requires reconciliation of conflicting constraints and trade-offs. Here, the (+) strand must fold (and remain folded) into a catalytic RNA motif and interact with the complementary (-) strand template without forming the thermodynamically highly-favored (+)(-) strand duplex, which is an inactive, dead-end product. Again, we could resolve this conflict by exploiting the ability of triplets to shift the folding and duplex-formation equilibrium to kinetically trap the QT45 (-) template in an extended template conformation (fig. S13, (*32*)).

One might assume that simply providing an excess of QT45 (+) ribozyme over QT45 (-) template should maximize synthetic yields. However, while this can improve the yields of partial products, we observed consistent inhibition of extension before reaching full-length QT45 (+) strand synthesis (fig. S15). Inhibition was even observed when supplementing the reaction with the unrelated 5TU polymerase ribozyme, pointing towards sequestration of the template (-) strands by the QT45 (+) strands independently of the catalyst used. As (+) strand outcompetes triplet substrates for binding to the (-) strand template, it drives increasing formation of the unproductive (+)(-) strand duplex (fig. S13). Indeed, in the absence of the QT45 (+) strand, we observed efficient full-length synthesis by 5TU. This suggests that the strand-inhibition problem, caused by the formation of QT45 (+)(-) strand duplex, is a key obstacle to closing the self-replication cycle.

We hypothesized that substrates that interact more strongly with the template strand might compete better with QT45 (+) for hybridization to the QT45 (-) template. To test this, we performed a series of reactions each supplemented with a single triphosphorylated RNA hexamer substrate complementary to different template positions. Among these, hexamers binding to the template region for the AU-rich apical loop (G25-C30, or A22-U27) proved effective at relieving strand-inhibition and supporting full-length synthesis (fig. S16, Fig. 4B). Therefore, the inclusion of one defined RNA hexamer together with the required triplet substrates enabled self-synthesis of the full length QT45 (+) strand by the QT45 (+) ribozyme on a QT45 (-) template (Fig. 4B). Deep sequencing confirmed the synthesis of the correct QT45 (+) product sequence (Fig. 4C). This shows that the QT45 ribozyme can complete the individual steps of a self-replication cycle in two separate reactions, using a random pool of trinucleotide substrates for the synthesis of its complementary strand, but needing defined triplets and one hexanucleotide to complete the synthesis of its self strand.

Using a random pool of all possible trinucleotide substrates enables the introduction of mutations during replication, which is a prerequisite for evolution. However, the number of mutations per round of replication must be bound within the “error threshold” (*6, 9, 42*). The error threshold is determined by the fidelity of replication and the relative replication fitness of the most active sequence and its mutants, and thereby limits the maximum sequence length that can be replicated without loss of information. The short length of QT45 substantially reduces its error threshold. If the fidelity observed for the QT45 synthesis of its complementary (-) strand (94.1%) persisted in a full self-replication cycle, it would be sufficient to maintain its own genome even at a modest relative fitness (fig. S17).

We also observed a low background of full-length product formation independent of triplet substrates, without the characteristic extension ladder of *bona fide* ribozyme synthesis products (fig. S1). This side reaction arises from recombination between partially extended products and the ribozyme through transesterification (fig. S18), via nonenzymatic ligation of a 2′-3′-cyclic phosphate to an adjacent 5′-hydroxyl as observed previously (*43–46*). In order to discern the synthetic products (produced by ribozyme catalysis) from the recombination products, we appended a short DNA tail to the QT45 ribozyme sequence, but not to the template encoding it (fig. S19) resulting in different migration of the synthetic (no tail) compared to the recombined (with tail) products in gel electrophoresis and allowing us to distinguish the two products in sequencing reactions. While recombination here is an inconvenient side reaction for analytical purposes, the inherent property of RNA to recombine might be advantageous in self-replication reactions as it would both accelerate prebiotic evolution and allow RNA to escape the deleterious effects of mutational drift (Muller’s ratchet) (*47–49*).

## Discussion

Our study shows that the complex functions needed for RNA replication can be performed by an RNA motif of just 45 nucleotides. While QT45 is a slow catalyst and replication is both low-yielding and requires separate reactions for (+) and (-) strand copying (and in the case of the plus strand reaches completion only with the aid of a hexamer substrate), it is encouraging that QT45 can perform the basic functions of RNA self-replication with a fidelity potentially sufficient to sustain its short genome. Furthermore, this ribozyme has only undergone a total of 18 rounds of evolution from a random sequence pool, underlining a likely potential for further development.

Two key conflicting constraints are inherent to RNA self-replication: the simultaneous coexistence of ribozymes as both folded catalysts and unfolded templates, and the inhibition of the ribozyme catalyst by hybridization to its complementary template strand. The QT ribozyme-catalyzed synthesis of itself and of its complementary strand shows how these constraints can be reconciled via the emergent properties of RNA ensemble equilibria and their modulation by substrate interactions and reaction conditions. Together with recent advances in our understanding of the physico-chemical conditions conducive to RNA replication (*32, 50–52*) this may enable the establishment of iterative cycles of RNA-catalyzed RNA self-replication.

The small size of the QT ribozymes not only reduces the synthetic and fidelity burden for self-replication but suggests that motifs encoding this activity are likely more abundant in RNA sequence space than anticipated. This substantially increases their accessibility by nonenzymatic RNA polymerization (*25, 26*), narrowing the conceptual gap between enzymatic RNA replication and prebiotic chemistry, and thus increasing the plausibility of RNA-based replication at the origin of life.

## Acknowledgments

We thank Luca Schwarz for triphosphorylated hexanucleotide purification and validation, Dr. Ben Porebski for advice with programming, Dr. Kim Liu for advice on sequencing, Dr. Isaac Gallego and Dr. Niklas Freund for T7 and TGK polymerase expression and purification, and Dr. Enrico S. Colizzi for useful discussions.

## Funding

The research at MRC LMB was supported by the Medical Research Council, as part of United Kingdom Research and Innovation (also known as UK Research and Innovation (UKRI)) [MC_U105178804] (EG, CJKW, JA, SLYK, KG, BEC, PH), a grant from the Volkswagen Foundation [96 755] (EG, BEC), a Herchel Smith studentship (2017) (CJKW) and a Cambridge trust PhD fellowship (SLYK). For the purposes of open access, the MRC Laboratory of Molecular Biology has applied a CC BY public copyright license to any Author Accepted Manuscript version arising.

## Author contributions

EG, JA and PH conceptualized and supervised the project, EG carried out all the work described except library construct design (with JA, CJKW), fidelity analysis (with SLYK), fitness landscape mapping (SLYK, CJKW with EG), 2’-hydroxyl substitutions mapping (KG with EG), catalytic rate (BEC). EG and PH wrote the manuscript with inputs from all authors.

## Competing interests

authors declare that they have no competing interest.

## Data and materials availability

sequencing reads and code to analyze them will be made available on Dryad and Zenodo.

## Notes

### Competing Interest Statement

The authors have declared no competing interest.

