## Supplementary Materials for "A polymerase ribozyme that can synthesize both itself and its complementary strand"

**The PDF file includes:**

Materials and Methods

Figs. S1 to S19

Tables S1 to S3

### Materials and Methods

1. Various nucleic acid manipulations

Methods repeatedly used in the paper have been collated for convenience. Variations from standard protocols are otherwise stated *in itinere.*

#### 1.1. T7 *in vitro* transcription

The *in vitro* transcription method used is based on (*53*). If the RNA required a triphosphate at the 5′-end, the "GTP" transcription protocol was used. If the RNA required a monophosphate at the 5′-end, the "GMP" transcription protocol was used. "GTP" transcription reaction conditions: 40 mM Tris-Cl pH 8, 10 mM DTT, 2 mM spermidine, 20 mM $MgCl_2$, 7.5 mM each NTP (Thermo Fisher Scientific), double-stranded DNA template containing 5T7 sequence at the 5′ end upstream of the region to transcribe (varying amount, preferably >5 pmoles), 0.01 units/µL of inorganic pyrophosphatase (Thermo Fisher Scientific), ~50 µg/mL of T7 RNA polymerase (expressed and purified in house). Reactions were incubated overnight (~16 hours) at 37°C. In order to remove template DNA, reactions were treated with 0.1 units/µL of Turbo DNase (Invitrogen) for 1 hour prior to purification. "GMP" transcription reaction conditions varied the nucleotide concentration as follows: 4mM each NTP, 20mM GMP. All other components were not varied from the "GTP" transcription.

#### 1.2. PAGE purification of oligonucleotides

Oligonucleotides derived from *in vitro* transcription or chemical synthesis was mixed in in FA9525 denaturing loading buffer (95% formamide, 25 mM EDTA, bromophenol blue) to a final concentration of >60% formamide. Samples were heated at 94°C for 5 minutes to denature the nucleic acid and separated on an 8 M Urea 1xTBE denaturing PAGE. Gels were run at constant 30W on an EV200 Large Format PAGE Unit Gel Unit (Cambridge Electrophoresis) using bromophenol blue as a marker for migration. UV shadowing was used to identify the band of interest. The gel fragment containing the band of interest was excised, crushed using a pipette tip, and suspended in TE buffer (10mM Tris-HCl pH 7.4, 1mM EDTA). The slurry was frozen in dry ice, thawed at 50 °C for 5 minutes and left rotating at 4 °C (2 hours to overnight). The eluate was filtered using a Spin-X 0.22 $\mu$m cellulose acetate filter (Costar) and precipitated in 73% ethanol (for ribozymes/long oligonucleotides) or 85% ethanol (oligonucleotides < 8 nt).

Absorbance at 260nm of the purified nucleic acids was measured using a Nanodrop ND-1000 spectrophotometer (Thermo Fisher Scientific) and the concentration was determined based on the measured absorbance and the sequence using Oligocalc (*54*)

### 1.3. TGK RNA synthesis

TGK (a primer dependent DNA-dependent RNA polymerase protein using single-stranded DNA as template (*55*)) was used to synthesize marker oligonucleotides. RNA primers as in the reaction requiring a marker were used. The primer extension reactions were carried out in 1x Thermopol buffer (NEB), 3 mM $MgSO_4$, 0.625 mM each NTP, 0.5 µM primer, 1 µM template, 150 nM TGK (94 °C 10 seconds, 40 °C 1 minute, 65 °C 1 hour, repeated once).

### 1.4. Adenylation

5′-end phosphorylated, 3′-end blocked DNA adapter was incubated at 20 µM for 2h at 65°C in 1x 5′ DNA Adenylation Reaction Buffer (NEB) supplemented with 1 mM ATP (Thermo Fisher Scientific) and 5 µM of Mth RNA ligase (NEB). Reaction size was usually 80 µL, but it was scaled depending on need. DNA from the reaction was then PAGE purified alongside a non-adenylated DNA adapter that serves as control marker.

### 1.5. Ligation of RNA to pre-adenylated adapter

T4 RNA ligase 2 truncated KQ (NEB) was used to ligate the 3′-end of single-stranded RNA to a DNA adapter without a bridging oligonucleotide. Each reaction contained a maximum of 1µg/µL RNA-bound Dynabeads MyOne Streptavidin C1 (ThermoFisher Scientific) beads or 50 nM of RNA in solution. Ligation was carried out in 1x NEB RNA ligase buffer, 15% PEG8000, 2 µM adenylated adapter, 0.04% Tween 20 and 20 U/µL of ligase for 2 hours at room temperature.

### 1.6. Template preparation for *in vitro* transcription

The indicated DNA oligonucleotides were cross-extended in four cycles of thermal cycling in standard reaction conditions for GoTaq HotStart Green MasterMix (Promega). Reactions were purified using Qiaquick PCR purification kit (Qiagen) for products >100 base pairs, and with a Nucleotide Removal Kit (Qiagen) for products <100 base pairs in length.

#### 1.7. Triplet and dimer transcription

Triplets and dimers were prepared via run-off *in vitro* transcription using T7 RNA polymerase. A detailed method can be found in (*56*). Briefly, reaction conditions were varied as follows: 100 pmoles of template for each triplet was mixed with equimolar 5T7. The "GTP" transcription protocol with the exception that a lower total NTP concentration was used (4.32 mM) as this yielded better defined bands for purification. 50 µL transcription reactions were stopped by adding 2 µL EDTA and 5 µL of 100% glycerol. The reaction products were separated on a 30% 3 M Urea denaturing PAGE gel using an EV400 DNA Sequencing Unit (Cambridge Electrophoresis). UV shadowing was used to identify bands, and the correct band, identified based on relative migration to known triplets was excised. The RNA was extracted from the gel fragment as in method 1.2. Correct sequence composition was confirmed by A260/280 absorbance ratio, measured with a Nanodrop ND-1000 spectrophotometer (Thermo Fisher Scientific)

#### 1.8. Template-dependent ribozyme-catalyzed RNA synthesis: reaction setup-up and product detection

##### 1.8.1. Reaction setup

Standard primer extensions reactions were typically conducted as follows: 5 pmoles of biotinylated fluorescently labelled primer was mixed in water with template, ribozyme (5TU+t1.5 or QT), triphosphorylated oligonucleotide substrate(s), and annealed (80°C 2 minutes, 17°C 10 minutes) in half the final reaction volume. The reaction components were then kept on ice until pre-chilled buffer and salts were added. Final buffer concentrations are described in each figure legend. Upon buffer addition the reactions were equalized on ice for ~1 minute, then frozen in dry ice for ~ 1 minute and transferred to a -7°C R4 series TC120 refrigerated cooling bath (Grant) for the time indicated in figure legends. Reactions were then stopped with equimolar EDTA to the $Mg^{2+}$ in the reaction.

The concentrations indicated in the figure legends describe pre-freezing conditions, assuming that the final post-freezing operational volume is determined by the solute concentration in the sample. The formation of ice-crystals causes all solutes to concentrate to their final operating concentrations upon equilibration of the eutectic phase. pH is also expected to vary due to the temperature dependence of the buffer's pKa (*57*). For example, if 1 µmole of KCl and 1 pmole of RNA were added to a reaction volume of 10 µL or 20 µL, both are described as 100 mM KCl and 0.1 µM RNA of a virtual 10 µL pre-freezing reaction volume, as the concentrations in the eutectic phase would be identical post-freezing.

#### 1.8.2. Product detection for analytical purposes

For biotinylated primers, reactions were incubated with Dynabeads MyOne Streptavidin C1 (Thermo Fisher Scientific) at 0.1 pmol biotinylated primer/µg of beads, in at least one reaction volume of BWBT (0.2 M NaCl, 10 mM Tris-HCl pH 7.4, 1 mM EDTA, 0.1% Tween-20). Beads were washed once in BWBT, twice in NaBET25 (25 mM NaOH, 1mM EDTA, 0.05% Tween 20), and again washed twice in BWBT before resuspending in FA9525 and heated at 94°C for 5 minutes in order to disrupt the biotin-streptavidin interaction and elute the primer from beads. The supernatant was run on an 8 M urea 1x TBE denaturing PAGE. Gels were analyzed on a Typhoon Trio scanner (GE Healthcare). For non-biotinylated primers, reactions were mixed in >60% FA9525 with 10-20-fold molar excess of unlabeled strand complementary to the template in the reaction (described as "competing oligos" in table S3) to prevent product/template reannealing. Upon denaturation (94°C 5 minutes) RNAs were separated on a 8 M Urea 1xTBE denaturing PAGE. Gels were analyzed on a Typhoon Trio scanner (GE Healthcare). Gel bands were quantified using the ImageQuant analysis software.

#### 1.8.3. Product detection and recovery for selection and sequencing

Dynabeads MyOne Streptavidin C1 were added to the stopped reactions at 0.05 pmol biotinylated primer/µg of beads or an even greater excess of beads, in at least one reaction volume of BWBT. Then, the beads were washed twice with BWBT, twice with NaBET25 to confirm covalent linkage of construct to primer (and transferred to a fresh microcentrifuge tube to minimize downstream contamination between washes), and twice with BWBT, before resuspending in FA9525. Biotinylated RNA was eluted from the beads by disrupting the biotin-streptavidin interaction by heating at 94 °C for 5 minutes, and separated on a 8M Urea 1X TBE denaturing PAGE gel alongside RNA markers equivalent to successfully ligated constructs (generated by similar extension reactions but with added 5TU and t1.5 ribozymes). The marker-adjacent gel region in the construct lane was excised. Biotinylated RNA was then eluted and bound to MyOne Streptavidin C1 Dynabeads in BWBT overnight. After 50 µm filtering (Partec Celltrics(Wolflabs (York, UK))) of the supernatant to remove gel fragments, the beads were washed twice with BWBT, twice with NaBET25 (transferred to fresh tube between washed), and twice with BWBT before proceeding to subsequent reactions.

### 2. De novo selection

#### 2.1. Dimeric construct selection protocol

Details specific to each selection round can be found in table S1. A diagram of the steps involved in the dimeric selection construct can be found in fig. S1. Round 1 constructs were prepared via the circularization of a starting materials of 200 pmoles of ULTc2dN40/ULTc2dN30/ULTc2dN20 using equimolar 5T76FfGG as a splint. The two oligonucleotides were annealed (80 °C 2 minutes, 17°C 10 minutes) in 4/5 of the final reaction volume in 1X T4 DNA ligase reaction buffer (NEB). The components were kept on ice until prechilled T4 DNA ligase diluted in 1/5 of the reaction buffer was added at a final concentration of 8000 units/µL. Each oligonucleotide in the reaction was at a low concentration of 100 nM to favor intramolecular ligation. After 1 h incubation at 16 °C, dNTPs (GE Healthcare UK) were added to a final concentration of 800 µM and T4 DNA polymerase (NEB) was added at a final concentration of 0.012 units/µL. These were incubated at 25 °C for an additional hour. The product was purified using Qiaquick PCR Purification kit (QIAGEN).

From round 2 onwards, the libraries required an additional reaction step for the generation of a ssDNA equivalent to ULTc2dN40/ULTc2dN30/ULTc2dN20. To do so, 5 µL of RT-PCR reaction from the previous round was used as template for PCR with GoTaq HotStart Green MasterMix (Promega) for 14 cycles. 200 pmoles of primers forceGG17 and AACA2ULT were used. DNA Polymerase I, Large (Klenow) fragment (NEB) was added after PCR at a final concentration of 0.05 units/µL and incubated at room temperature for 15 minutes to generate blunt ends. Products were purified using a Nucleotide removal kit (QIAGEN). Lambda exonuclease (NEB) was used to remove the phosphorylated strand and generate ssDNA for subsequent steps, and repurified. The ssDNA was annealed with equimolar 5T76FfGG as a splint, and treated similarly to round 1, with an additional 5′-end phosphorylation step prior to ligation (1X T4 DNA ligase buffer, 0.25 units/µL T4 PNK (NEB), 30 minutes at 37°C, heat inactivated 20 minutes at 65 °C). The same reaction, diluted 5 fold in the same ligation/extension buffer described for round 1 except for a lower T4 DNA ligase concentration of 1600 units/µL. The product was Qiaquick PCR purified (QIAGEN).

After obtaining this purified product, 3 further reactions were carried out sequentially in one pot. The first reaction consisted in nicking of deoxy-uracil nucleotides in the oligonucleotide via USER enzyme treatment (0.05 units/µL USER enzyme (NEB) in 1X Cutsmart buffer supplemented with 5 mM DTT for 15 minutes at 37 °C), the second reaction was the dephosphorylation of 3′-ends (T4 PNK (NEB) at 0.5 units/µL for 45 minutes at 37 °C), lastly dNTPs and DNA Polymerase I, Large (Klenow) fragment (NEB) at a final concentration of 1 mM and 0.05 units/µL respectively were added to extend the 3′-ends, and incubated for an additional 45 minutes at 37 °C. Reactions were stopped using 6X Purple Gel Loading Dye (NEB) and products were separated on a 1X SYBR Safe (Life Technologies Ltd) 3.5% UltraPure Agarose (Life Technologies Ltd) gel. The correct products were excised, gel purified (QIAGEN) and used as templates for “GTP” transcription. The transcription were treated with Turbo DNase (Invitrogen) and PAGE purified as in method 1.2.

Reactions were set-up as in method 1.8 with equimolar primer/template to the RNA libraries (exact conditions for each round in table S1). The final concentrations before freezing were 50 mM $MgCl_2$, 200 mM KCl, 50 mM CHES-KOH pH 9, 0.05% Tween 20, with 50 nM primer/template/library in round 1 and round 12, and 20 nM in all other rounds. Reactions were stopped with equimolar EDTA to the $Mg^{2+}$ present and products were recovered and bound to beads as described in section 1.8.3. The beads were then resuspended in the RT-PCR reaction containing pTLT reverse primer and varying forward primer covering part of the ligation junction from the ribozyme reaction (details of primers used in each round of the selection are in table S1). The RT-PCR product DNA was used in subsequent selection rounds or sequenced. Error prone PCR was carried out after round 5 using the GeneMorph II kit for mutagenesis (Agilent).

### 2.2. Monomeric selection

The selection in monomeric form followed similar steps to the one in dimeric form, with a simpler construct generation procedure. The ribozyme primer extension reaction set-up was identical, with recovery only diverging just before RT-PCR. The bead-bound biotinylated RNA was 3′-end dephosphorylated using T4 PNK and adaptor ligated to a pre-adenylated adapter (HDVlig) prior to RT-PCR using primers HDVrec and forceGG (or a variable primer depending on the template used in the ribozyme primer extension, detailed in table S3). The RT-PCR product was then used as a template for a subsequent in-nest PCR, with primers 5T76FfGG and HDVrt used regenerate the T7 promoter and a HDV ribozyme 3′-end cassette. The DNA was subsequently purified using Qiaquick PCR purification kit (QIAGEN) and used for “GTP” transcription. The transcription was treated with Turbo DNase (Invitrogen) and PAGE purified as in method 1.2. Upon selection (which followed the same steps as the dimeric selection of method 1.2), the recovered bead-bound RNA was dephosphorylated with T4 PNK for 1 hour, adapter ligated to AdeHDVlig as in method 1.5. The product was amplified via RT-PCR with the primers specified in table S3 using SuperScriptIII/Platinum Taq One Step RT-PCR system (Thermo Fisher Scientific). The resulting product was used in subsequent selection rounds or sequenced.

### 3. Regiospecificity assay

The ribozyme clones from Fig. 1C were used to carry out a primer extension reaction from a primer containing a single G at the 3′-end (FITCrec3), opposite a template (t6Freg3GCG2CUG) encoding 2 GCG triplets and 1 CUG. Reactions were carried out using 20 pmoles of equimolar primer, template, and ribozyme in a final concentration of 50nM. 5 µM each pppGCG and 5′-HO-CUG triplets were used, and the reactions were incubated for 4 days at -7 °C frozen. The full-length extension product was PAGE purified, precipitated, and incubated with 50 units of RNAse T1 in 2 mM EDTA, 50mM Tris-HCl pH 7.4 for 20 minutes at 37 °C. The reaction was analyzed on an 8 M Urea 1X TBE denaturing PAGE, using FITCreg3P as a marker for cleaved reactions.

### 4. Hammerhead ribozyme activity assay and sequencing

The hammerhead synthesis reaction conditions were described in the figure legend. Three different primers were used depending on the subsequent steps. For visualization, the reaction was set-up using BCy3P10, to assay the activity of the synthesized product, A647BP10 was used in order to simultaneously scan the synthesized hammerhead and its FAM labelled substrate. For sequencing, a primer containing a 5′ DNA overhang was used (bioCy3KyleP10) to facilitate product amplification. Synthesized products were recovered as in method 1.8.3 and used for the subsequent step.

To assay hammerhead ribozyme activity, a two-fold molar excess of FAM labelled substrate (Fsubuhl) was mixed with A647 labelled hammerhead (either synthesized by QT51, 5TU or TGK). Reactions contained 10 mM Tris-HCl pH 8, 2 mM $MgCl_2$, and 0.05% Tween-20. Prior to $MgCl_2$ addition the reaction was incubated at 60 °C for 1 minute, then moved to ice. To start the reaction $MgCl_2$ was added and the sample was frozen in dry ice, and incubated at -7 °C for 24 hours to let the reaction occur in the eutectic concentrate. The reaction was quenched in 65 % FA9525 and analyzed via denaturing PAGE as in method 1.8.2.

### 5. Ribozyme-catalyzed synthesis of itself and its complementary strand

For (+) strand self-synthesis reactions all reaction components (described in detail in the corresponding figure legend) were mixed in a total volume of 125 µL at room temperature. To reduce recombination (as in fig. S18), the ribozyme used contained a 5′-phosphate (pQT45). To reduce strand-reannealing, a single cycle of heating and acidification followed by quick freezing was used to initiate the reactions (as in (*32*)). Reactions were then incubated at -7 °C for the time indicated in the figure legend.

For (-) strand synthesis, the reaction was set-up as in method 1.8, with the conditions described in the figure legend.

### 6. High-throughput sequencing

#### 6.1. Sequencing libraries preparation

Libraries for Illumina sequencing were prepared from previously amplified RT-PCR product, by further PCR amplification with primers containing 5′ overhangs that introduce features needed for sequencing. The PCR was carried out using GoTaq HotStart Green MasterMix (Promega). An example of the PCR primers used to prepare libraries is shown below. Italicized is the region that hybridizes to the flow-cell. Underlined is the section where sequencing primers hybridize on the sequencer. A random three nucleotide “NNN” section is introduced to ensure high diversity in the beginning of the read. A four-nucleotide barcode sequence, denoted as “XXXX” is used to provide a unique barcode to each of the libraries sequenced in the same run, and it is de-multiplexed computationally after sequencing. “(land)” refers to the library-specific landing primer used.

>P5NNNXL
*AATGATACGGCGACCACCGAGATCTACAC*TCTTTCCCTACACGACGCTCTTCCGATCT**NNN**XXXX(land)
>P7NNNXL
*CAAGCAGAAGACGGCATACGAGAT*GTGACTGGAGTTCAGACGTGTGCTCTTCCGATCT**NNN**XXXX(land)

The amplified libraries containing the correct overhangs were agarose gel purified (Qiagen) and pooled together. Pooled libraries were quantified using a Qubit 2.0 Fluorometer (Thermo Fisher Scientific) using the high sensitivity dsDNA Quantification Assay Kits (Invitrogen), prior to denaturation and dilution in HT1 buffer (Illumina) to be sequenced on a MiSeq System, HiSeq 2500, or NextSeq 2000 (Illumina).

6.2. Analysis of sequencing data

We computationally pre-processed the sequencing data in order to facilitate downstream analyses. Paired-end reads were merged using PEAR (*58*). We then trimmed the single-end reads or merged paired-end reads using the BBtools pipeline (*59*) and quality filtered the reads for a minimum base quality of 30 over 100% of the read using the FASTX-Toolkit (*60*). The individual libraries were demultiplexed using Cutadapt (*61*).

For the fidelity analysis of ribozyme products, the pre-processed reads were aligned to the correct product using BBMap and Samtools. This generated a pileup file that was used for subsequent analysis and visualization using Python. The output pileup files were parsed in Python using the Pandas dependency and plotted as heatmaps using Matplotlib and Seaborn.  Alignments of a small number of clones were generated using MAFFT (*62*) on the Aliview visualization software (*63*).

### 7. Fitness landscape generation

#### 7.1. Selection library synthesis

For construction of the QT45 library for fitness landscape determination, equimolar amounts of oligonucleotides comprising different shifted registers of QT45 (QT45MO10, QT45MO10_iG1_dC45, QT45MO10_dG1_iC45, QT45MO10_C21D_ins1G, QT45MO10_C21D_ins45C, QT45MO10_C21D_ins20G, QT45MO10_C21D_ins22G, QT45MO10_U23D_ins45C, and QT45MO10_U23D_ins1G) were mixed together. In order to obtain oligonucleotides encoding the HDV ribozyme at the 3′ end, 2 nmol of the resultant mixture was then ligated to 2 nmol of HDVrest, using T4 DNA ligase and 2 nmol of HDVspl as a complementary DNA splint. After heat inactivation and spin concentration with a 3 kDa molecular weight cut-off filter, the construct was urea-PAGE purified. The product band was excised, eluted in 10 mM tris•HCl pH 7.4 overnight, and precipitated in 73% ethanol.

To generate the template for transcription, 260 pmol of the resultant oligonucleotide was then mixed with 260 pmol of partially-complementary oligonucleotide 5T76F6LfGG and Bst2.0 polymerase (NEB) was used to 'fill-in' the single-stranded regions to make a double-stranded product with a short linker (sp12-6L library template). Similarly, 20 pmol of resultant oligonucleotide from the above ligation step was also 'filled-in' with 5T76F8LfGG using Bst2.0 to make a similar product but with a longer linker (sp12-8L library template). Both products were then Qiaquick PCR purified (Qiagen) and transcribed overnight using T7 RNA polymerase to yield RNA selection libraries. Products were subsequently purified using preparative-scale urea-PAGE. The product band was excised, eluted in 10 mM tris•HCl pH 7.4 overnight, and precipitated in 73% ethanol.

#### 7.2. In vitro evolution cycle, recovery, and sequencing

10 pmol of sp12-6L library was annealed with equimolar temp6FP10UGC3 template, BCy3P10 primer, and 1 nmol of UGC triplet in 250 µl of water with 0.1% tween (80 °C 2 min, 17 °C 10 min). The annealed mixture was then placed on ice, 250 µl of chilled 2x extension buffer (100 mM CHES, 100 mM $MgCl_2$) was added, and the reaction was then frozen with dry ice and incubated at -7 °C for 7 hours.

After the incubation, the reaction was stopped with equimolar EDTA and products were recovered and bound to beads as described in section 1.8.3. Then, the 3′ end of the bead-bound constructs were first dephosphorylated with T4 PNK for 1 hour (with T4 PNK added after bead resuspension in other reaction components including 0.05% tween-20) and AdeHDVlig was subsequently ligated to the dephosphorylated 3′ end for 2 hours (with RNA ligase 2 truncated KQ added after bead resuspension in other reaction components including 0.4% tween-20). Beads were washed twice in BWBT and used in a 50 µl RT-PCR. Reverse transcription and PCR were carried out using HDVrec and P10UGCugFrec primers using the SuperScriptIII/Platinum Taq One Step RT-PCR system (Thermo Fisher Scientific). 4 µl of the resulting product was run on a 4% agarose gel to check the size of the products. The remaining product was Qiaquick PCR purified and then further amplified using primers that introduce indexed adapters for Illumina sequencing (P71forceGG_2024 and P51HDVba_2021) to generate the sequencing construct in a 50 µl GoTaq HotStart (Promega) PCR. Products were purified by 4% agarose gel, quantified on a Qubit 2.0 fluorometer, and then sequenced on a NextSeq 2000 (Illumina).

The above process was repeated to screen for activity on two other templates encoding 3 AUA and 12 CUA with similar setups. Regarding the 3 AUA extension setup, 10 pmol of sp12-6L library was annealed with equimolar temp6FnewnewP12AUA3 template, BCy3newnewP12 primer, and 2.5 nmol of AUA triplet in 250 µl of water with 0.1% tween (80 °C 2 min, 17 °C 10 min). The annealed mixture was then placed on ice, 250 µl of chilled 2x extension buffer (100 mM CHES, 100 mM $MgCl_2$) was added, and the reaction was then frozen with dry ice and incubated at -7 °C for 17 hours. Regarding the 12 CUA extension setup, 10 pmol of sp12-8L library was annealed with equimolar temp6FP10CUA12 template, BCy3P10 primer, and 2.5 nmol of CUA triplet in 250 µl of water with 0.1% tween (80 °C 2 min, 17 °C 10 min). The annealed mixture was then placed on ice, 250 µl of chilled 2x extension buffer (100 mM CHES, 100 mM $MgCl_2$) was added, and the reaction was then frozen with dry ice and incubated at -7 °C for 2 days. Downstream recovery processes were similar except that reverse transcription and PCR were carried out using HDVrec and newnewP12auaaFrec primers for the AUA sample, and HDVrec and forceGG primers for the CUA sample. Additionally, primers used to generate the sequencing constructs were also different. For the 3 AUA post-selection sample, primers P71forceGG_2024 and P53HDVba_2021 were used. For the 12 CUA post-selection sample, primers P71forceGG_2024 and P52HDVba_2021 were used.

Pre-selection libraries were also amplified using primers that introduce indexed adaptors for Illumina sequencing. The sp12-6L library was similarly PNK treated, adapter ligated with adeHDVlig, RT-PCR with forceGG and HDVrec, and finally PCR with P72forceGG_2024 and P512HDVba_2021. The sp12-8L library was processed similarly, except that P72forceGG_2024 and P514HDVba_2021 were used in the final PCR step.

7.3. Calculating fitness associated with each genotype

Reads from the NextSeq run were merged using PEAR(*64*) (where applicable) and reads from separate sequencing runs were combined. Reads were then demultiplexed into their respective libraries (input and 3 output libraries - 3 UGC, 3 AUA, 12 CUA) using Cutadapt, according to 6-nucleotide barcodes. Adapters were trimmed away also using Cutadapt, leaving only the variable QT45 sequence. Using FASTX-toolkit, reads were quality filtered such that each read contains only bases with Q-score 20 or above, and then identical sequences were collapsed while maintaining read count. Genotypes containing 10 reads or more in the input libraries, as well as at least 1 read in each of the 3 output libraries (3 UGC, 3 AUA, 12 CUA), were retained for downstream analysis to generate the single mutant fitness values. Genotypes containing 10 reads or more in the sp12-6L input library and at least 1 read in the 3 UGC output library were used to generate the double mutant fitness values. As the sub-library QT45MO10_dG1_iC45 was the most active and hence the most abundant in the output libraries, the fitness landscapes were generated using genotypes from this sub-library.

To determine the fitness associated with each genotype, the fraction of each library occupied by each genotype in the input and 3 output libraries were first calculated. The enrichment of each genotype during selection was then obtained by dividing the fractional abundance in the output library by that in the input library. Then, the fitness of each genotype was calculated as the log2 ratio of the enrichment of the genotype during selection, and the enrichment of the wild-type sequence during selection (*31*). Consequently, wild-type QT45 has a fitness of 0, while less active mutants have fitness values less than 0 and more active mutants have fitness values more than 0. An average of the three fitness values determined using the 3 output libraries was plotted to generate the single mutant fitness landscape, whereas the fitness value determined using the 3 UGC output library was used to plot the double mutant fitness landscape.

#### 7.4. QT39 fitness landscape

Determination of the QT39 fitness landscape followed a similar approach described above for QT45. However, instead of 3 output libraries of different templates encoding different triplets (3 UGC, 3 AUA, 12 CUA) used to generate the QT45 fitness landscape, 3 replicates of the same template (3 CUA) were used to generate the QT39 fitness landscape. Downstream processing also followed a similar procedure, with minor differences. Following PEAR merging of reads, reads were trimmed and demultiplexed with FASTX-toolkit, further trimmed with Cutadapt, then quality filtered to Q-score 30 or above and collapsed with FASTX-toolkit.

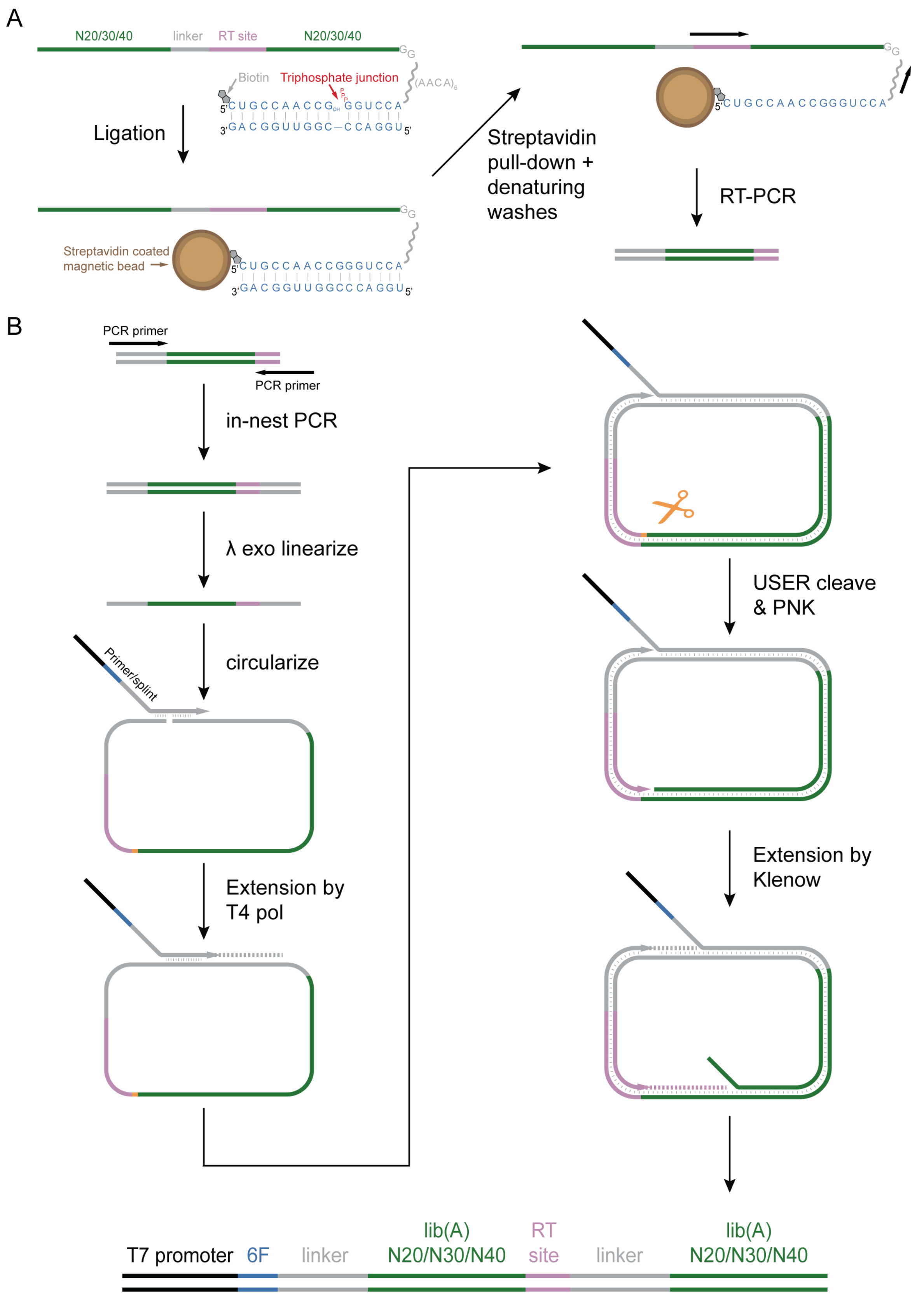


**Fig. S1 Homodimeric construct formation and selection scheme.**
(**A**) A homodimeric RNA library containing two copies of each random sequence is challenged to catalyze templated RNA ligation. Active members of the library are selectively recovered via streptavidin pulldown and a single monomer is amplified via RT-PCR. (**B**) The DNA of a single monomer is amplified via PCR and the product is linearized via λ exonuclease digestion. The linear product is circularized

using a DNA splint containing a 5'-overhang encoding the T7 promoter. The splint is then used as a primer to synthesize the complementary strand to the circular template. T4 DNA polymerase is used to avoid strand displacement and rolling circle amplification. A nick is formed in the circular strand using USER enzyme at a pre-defined position containing deoxy-uracil. After dephosphorylation of this nick, Klenow DNA polymerase is used to extend at the two available primer sites, displacing opposite strand, and generating the duplicated sequence of interest with a T7 promoter.

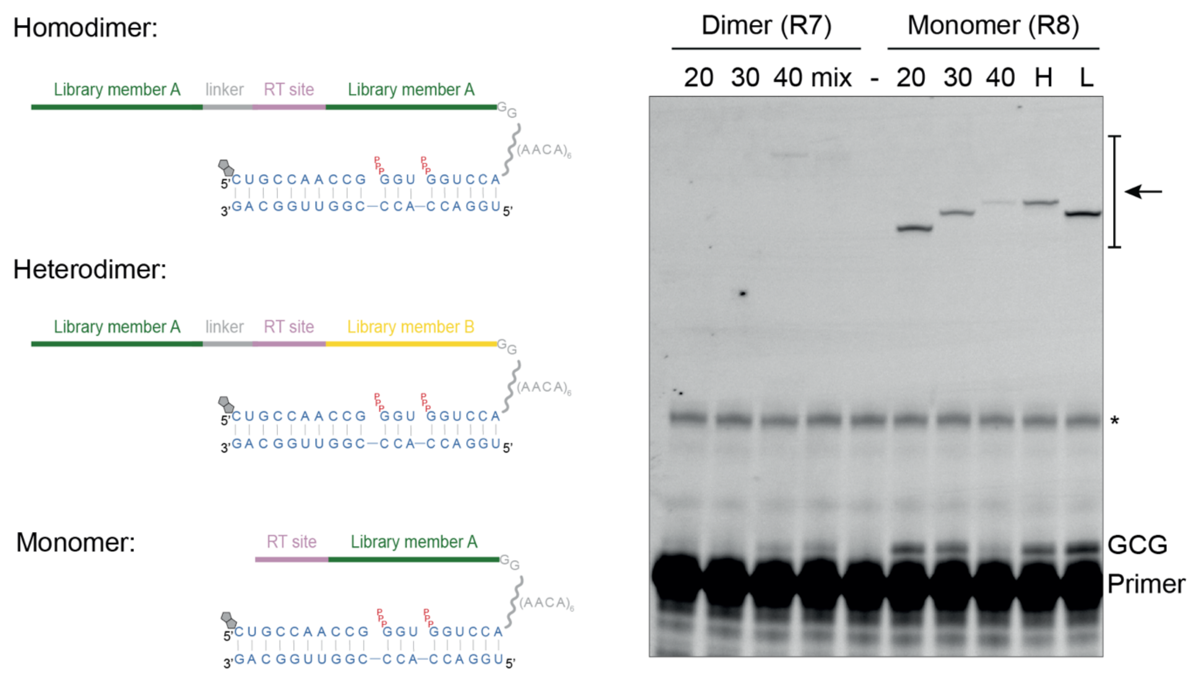


**Fig. S2: Library activity as monomer or dimer.**

(Left) Diagrams of three possible types of constructs are displayed. Homodimeric constructs contain two copies of the same library member. Heterodimeric constructs contain two library members differing in their sequence. Monomeric constructs have been truncated to only contain one library member and its RT site. (Right) Primer extension activity of the libraries as a dimeric construct in R7 compared with the libraries in monomeric construct in R8. Area where full length self-ligation is visible is indicated by an arrow. The asterisk indicates a reaction-independent anomalous primer migration band. Libraries of origin are annotated (see table S1 for more details). H/L consists in the “Mix” library’s individual components after purification. Reaction conditions: 0.5 µM primer BCy3newP10, 0.5 µM template temp6FnewP10GCG, 5 0.5 µM pppGCG, 0.05% Tween 20, 200 mM KCl, 50 mM $MgCl_2$, 50 mM CHES-KOH, pH 9, 16 hours at -7 °C frozen.

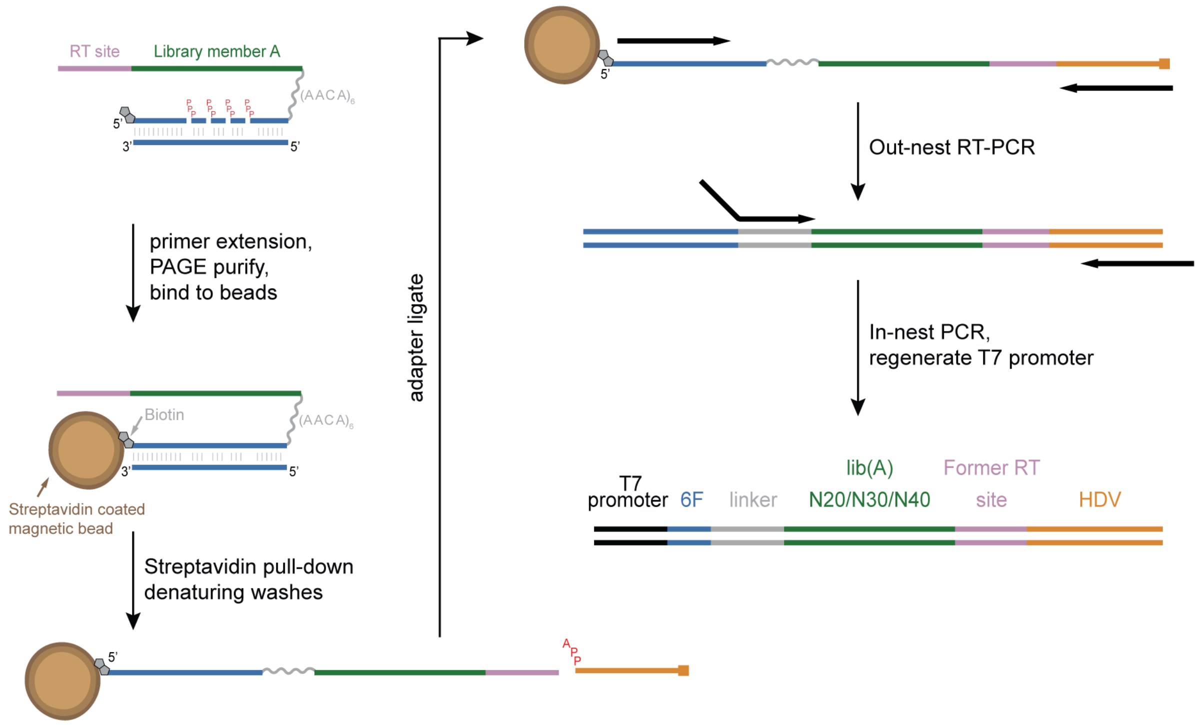


**Fig. S3: Selection in monomeric form.**

Diagram of the selection scheme used for the selection of a monomeric triplet polymerase ribozyme. An RNA construct containing the library of interest is incubated with a template, biotinylated primer and triplets. The 5' end of the selection construct contain 6 nucleotides that hybridize with the 5'-end of the template. Upon reaction, streptavidin beads are used to selectively recover the member of the library that carried out iterative triplet polymerization followed by ligation of the product to the hybridized region of the construct. Brief NaOH washes ensure recovery of solely the library members that are covalently bound to the biotin. Denaturing PAGE is used to select the correct size product. The recovered products are ligated to a DNA adapter, which provides a landing site for a primer in the subsequent reverse transcription. RT-PCR amplifies the recovered products, and a subsequent PCR is used to regenerate products suitable for transcription.

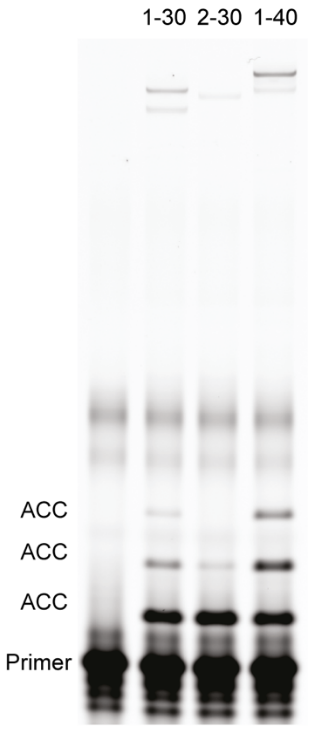


**Fig. S4: Polymerization of ACC.**
Iterative triplet polymerization by the clones displayed in (Fig. 1C) in the selection construct displayed in (Fig. 1B), with XXX=ACC and y=3. Reaction conditions: 50 nM ribozyme-substrate, 50 nM primer BCy3P10ga, 50 nM template temp6FP10gaACC3, 5 µM pppACC triplet, 0.05% Tween 20, 200 mM KCl, 50 mM $MgCl_2$, 50 mM CHES-KOH, pH 9, 3 days incubation at -7 °C frozen. Ribozymes are hybridized to the template.

A

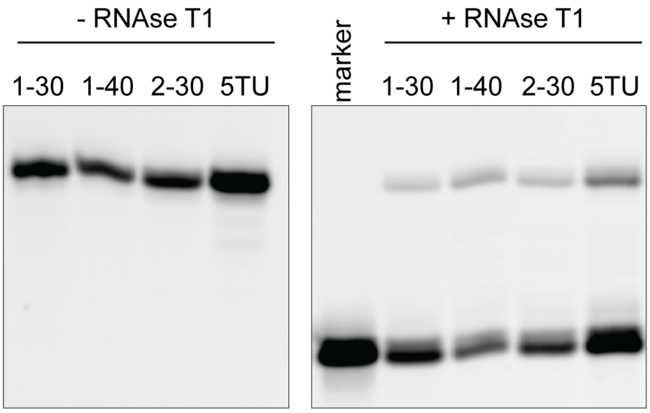


B

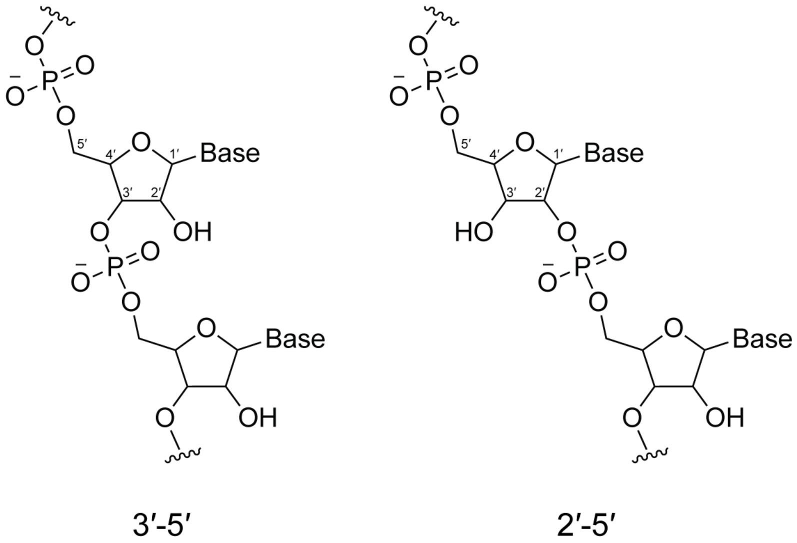


**Fig. S5: Regiospecificity of ligation assessed via RNAse T1 cleavage assay.**
(**A**) Ligation products by 1-30, 1-30, 1-40 and 5TU of a primer FITCreg3 containing a single G at its 3' end to a triphosphorylated substrate on template t6Freg3GCG2CUG are assayed for their susceptibility to RNAse T1 cleavage. All ribozymes tested generated a RNAse T1 (which cleaves 3'-5' linkage but not 2'-5') susceptible product, suggesting 3'-5' linkage. A 3' phosphorylated primer was used as a marker of correct digestion. (**B**) Chemical structure of the 3′-5′ phosphodiester bond and the 2′-5′ phosphodiester bond.

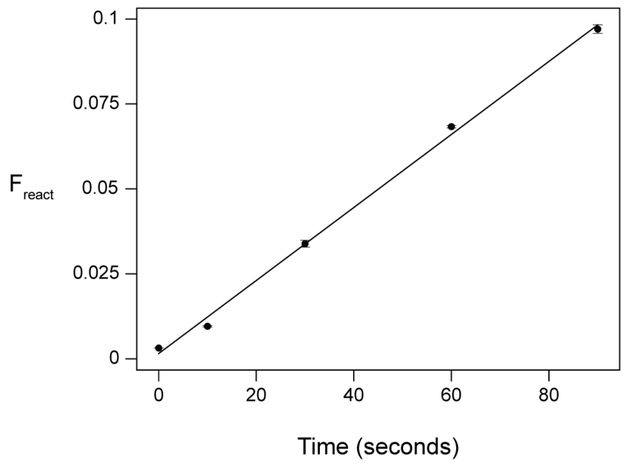


**Fig. S6: Catalytic activity of QT51**

Templated ligation of two RNA oligonucleotides in the format of Fig 1A, fraction of primer reacted is plotted over time in seconds. Reaction conditions: 0.1 µM ribozyme-substrate QT51_6F5L, 0.1 µM primer BCy3P10, 0.1 µM template temp6FF10, 500 mM $MgCl_2$, 250 mM CHES-KOH, pH 9, 0.05% Tween-20, 25 °C. Reaction were carried out as duplicates and error bars display the s.d.

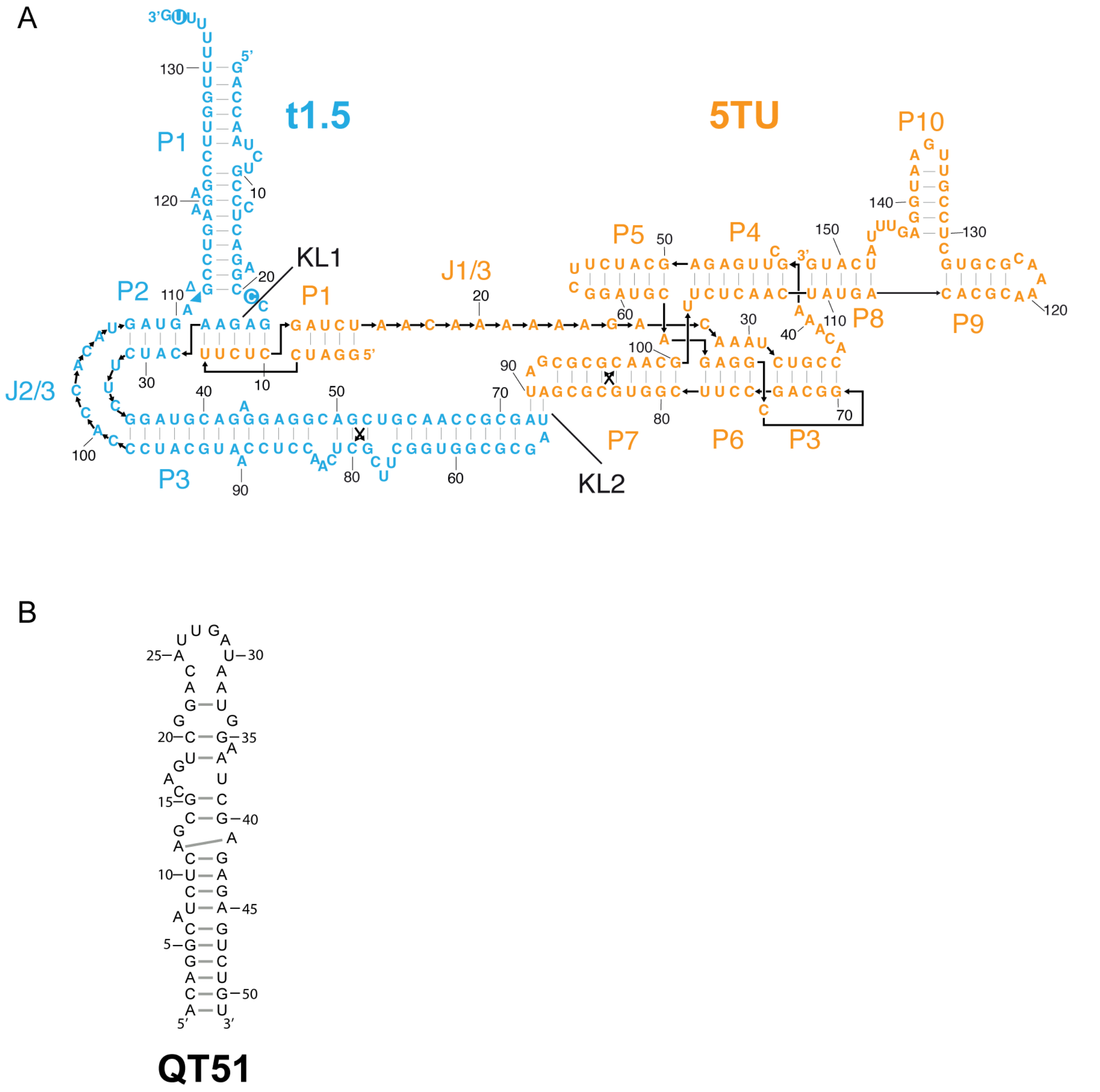


**Fig. S7 Comparison of secondary structure and size of 5TU+t1.5 and QT51 ribozymes.**
(**A**) 5TU ribozyme with t1.5 cofactor, mutations from the ancestral t1 sequence indicated on the diagram.
(**B**) QT45 ribozyme predicted secondary structure.

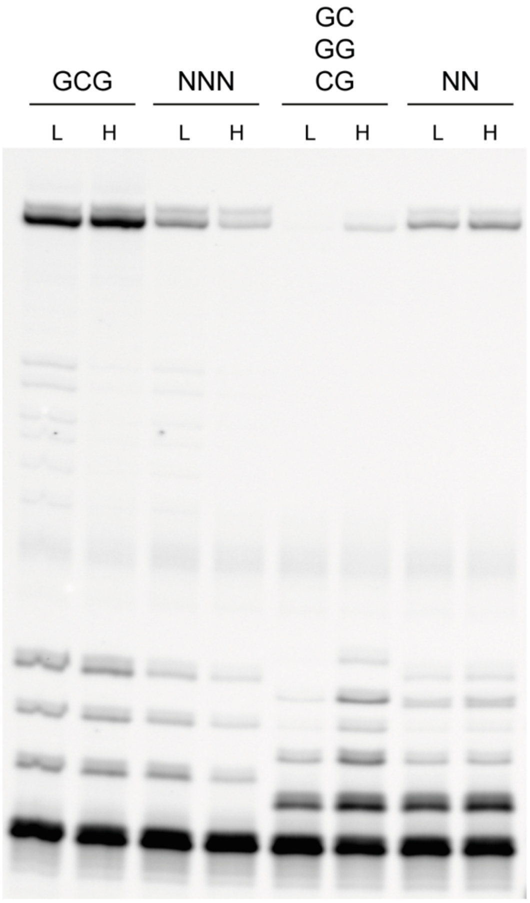


**Fig. S8: Promiscuity in substrate length.**

Templated polymerization of trinucleotides and dinucleotides in the format of Fig. 1B. Reaction conditions: 0.25 µM ribozyme-substrate 0_51_r7trim, 0.25 µM BCy3P10ga, 0.25 µM template temp6FP10gaGCG3, 50 mM $MgCl_2$, 200 mM KCl, 50 mM CHES-KOH, pH 9, 0.05% Tween-20, -7 °C frozen, 4 days. Triplet concentration L = 2.5 µM, H = 5 µM, Dinucleotide concentration L = 5 µM, H = 10 µM.

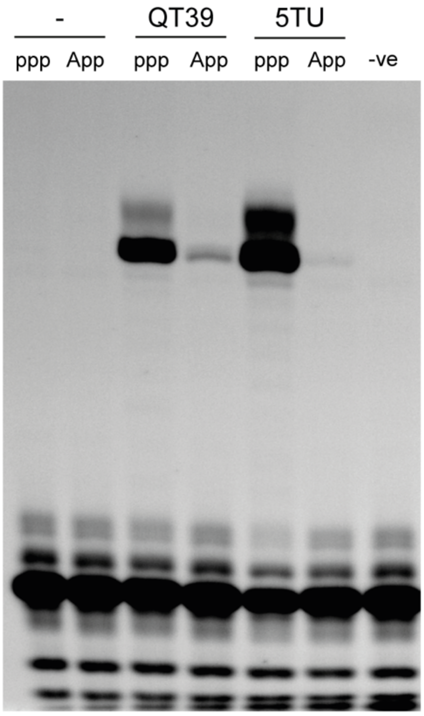


**Fig. S9: Promiscuity in substrate leaving group.**
Comparison of ligation activity between triphosphorylated and adenylated (App) downstream substrates. Reaction conditions: 0.25 µM primer F10, 0.25 µM template tempF10Ltest, 0.25 µM ribozyme (QT39a or 5TU+t1.5), 0.25 µM substrate (pppLtest1 or AppLtest1), 50 mM $MgCl_2$, 50 mM CHES-KOH, pH 9, -7 °C frozen, 19 hours.

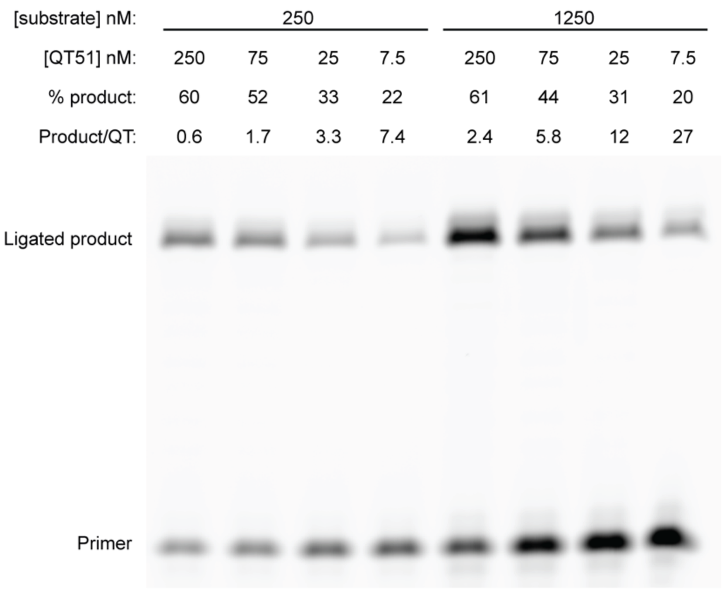


**Fig. S10: Multi-turnover activity.**
Ligation activity of QT51 over a range of ribozyme and substrate concentration. Ligated product per QT ribozyme is shown. Reaction conditions: 250 or 1250 nM primer/template/substrate (F10/tempF10Ltest/pppLtest1) as labelled, 250-7.5 nM QT51 ribozyme, 0.05% Tween-20, 50 mM $MgCl_2$, 50 mM CHES-KOH, pH 9, -7 °C frozen, 3 days.

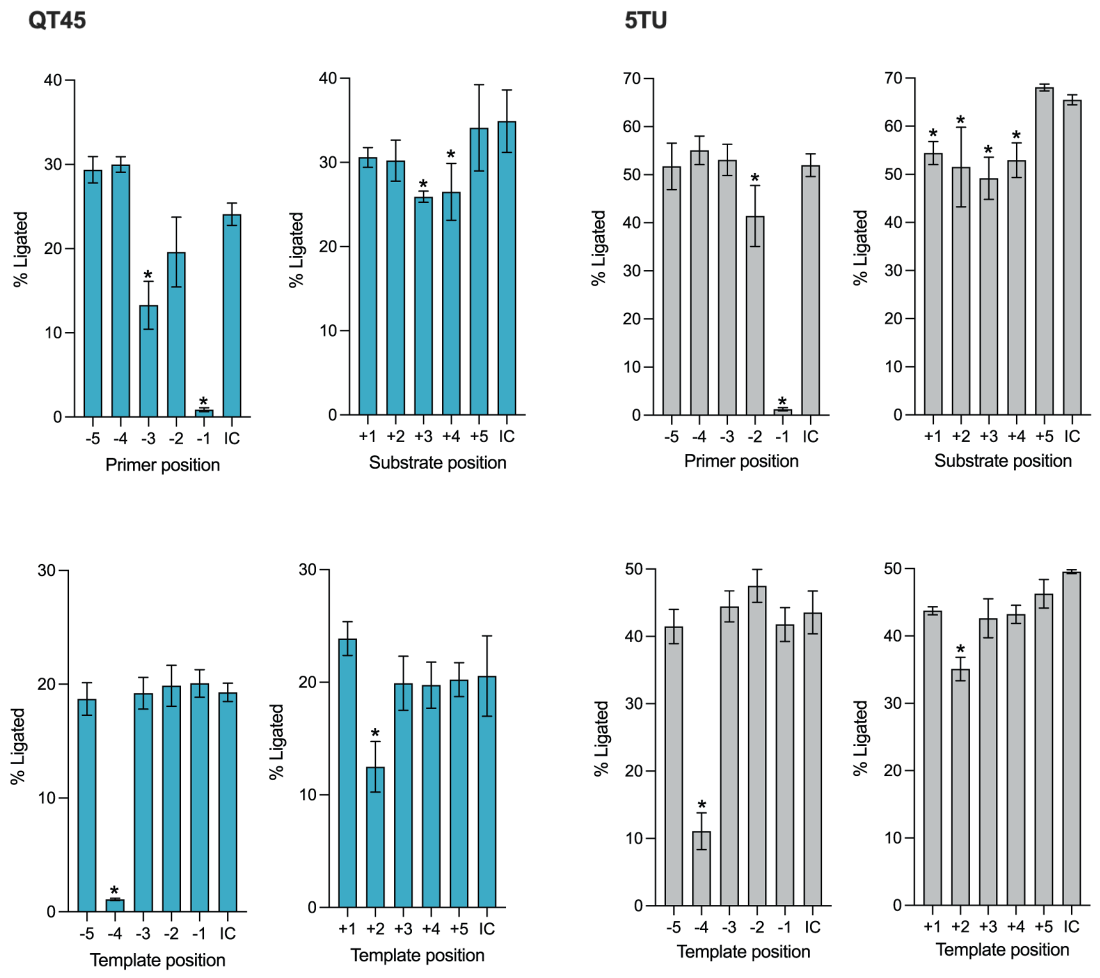


**Fig. S11 Effect of 2'-hydroxyl substitutions in the template, primer or substrate on ligation.** Percentage of product ligated by QT45 (teal) or 5TU+t1.5 (grey) in a single ligation junction where 2'-hydroxyl groups were individually substituted with 2'-deoxy, compared with an internal control (IC) of fully unsubstituted 2'-hydroxyl strand. Position of 2'-deoxy-substitutions indicated relative to a ligation junction (0). Reaction conditions: 0.1 µM primer F10, 0.1 µM template and substrate (see materials and methods for the exact sequences used), 0.1 µM ribozyme, 0.025% Tween-20, 50 mM $MgCl_2$, 50 mM CHES-KOH, pH 9, -7 °C frozen, 18 hours. Each substrate was tested in triplicates and the mean values for all positions were statistically analyzed using one-way ANOVA ($P < 0.05$) and a multiple comparison test to IC by the Dunnett Test; asterisks (*) mark the suppressed positions significantly different from IC.

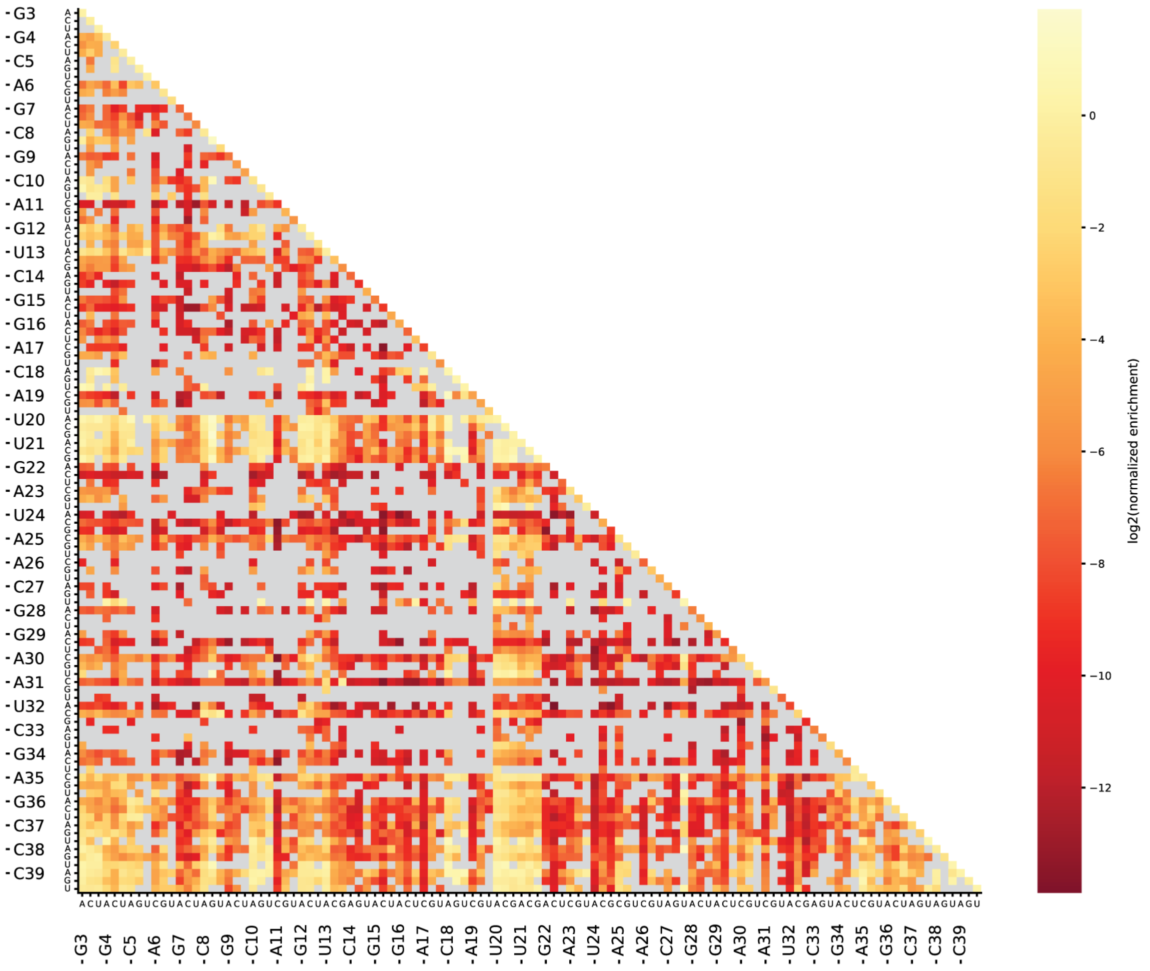


**Fig. S12 Heatmap for all measured single and double mutants of the QT39 ribozyme.** Heatmap for all measured single and double mutants of the QT39 ribozyme, with the first constituent point mutation indicated on the x-axis and the second one on the y-axis. Extension reactions on 3CUA template were done in triplicates and the mean fitness values were plotted. Fitness values are log-transformed enrichment values normalized to wildtype. Consequently, wild-type QT39 has a fitness of 0, while less active mutants have fitness values less than 0 and more active mutants have fitness values more than 0. As QT39 is a variant of QT45 that maintains the same core with a truncated stem, the QT39 heatmap shows similar trends as the QT45 heatmap (Fig. 2D).

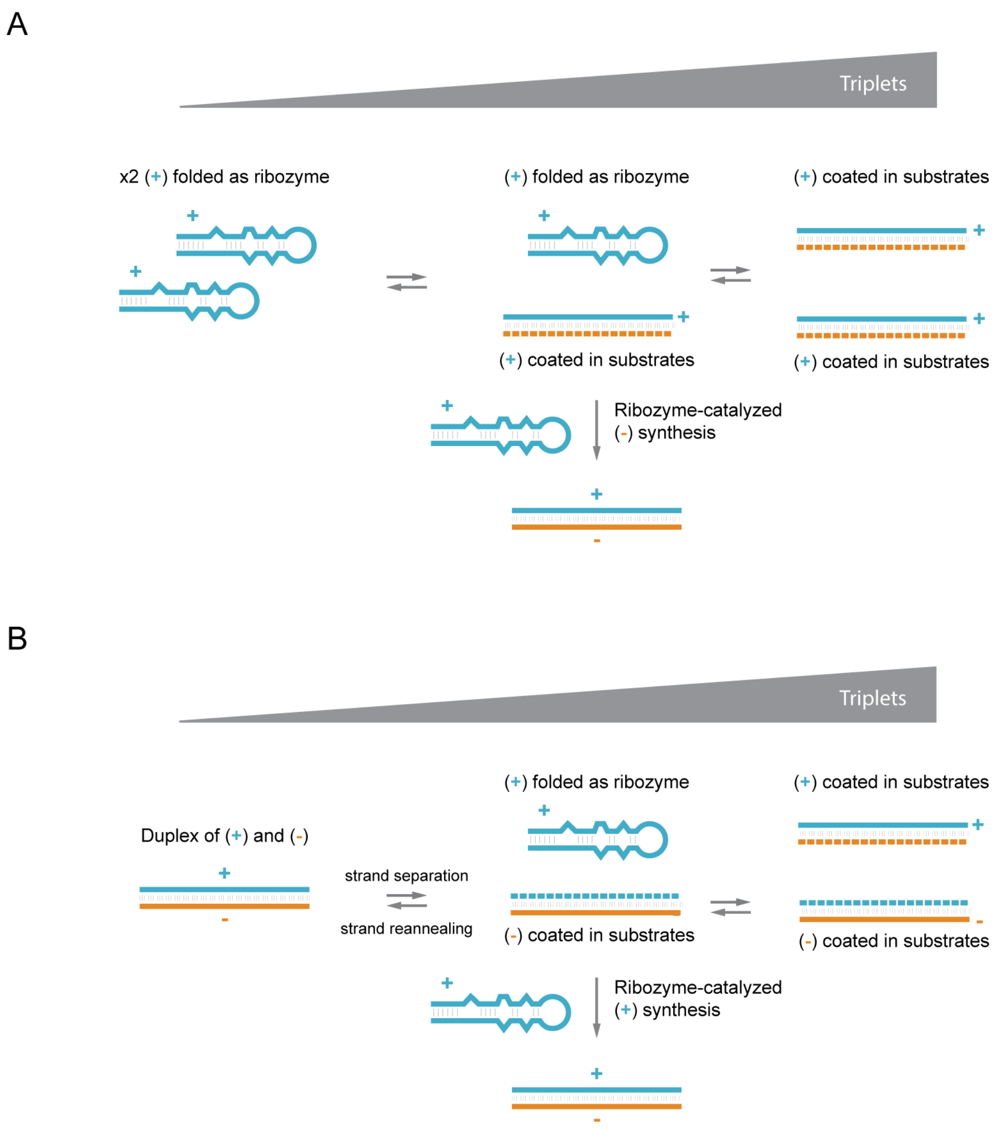


**Fig. S13: Proposed role of triplet concentration in RNA self-replication.**
(**A**) Effect of triplet concentration in (-) strand synthesis. At intermediate concentrations, triplet cooperatively unfold some ribozyme (+) strands, making them accessible as a template, while leaving other (+) strands folded as ribozymes. At higher concentrations all ribozymes are fully unfolded, and no (+) strand is available as catalyst. (**B**) Effect of triplet concentration on (+) strand synthesis. Upon strand separation at optimal triplet concentration, (+) strand folds as ribozyme and (-) strand is coated in substrates. At higher concentrations all ribozymes are fully unfolded, and no (+) strand is available as catalyst.

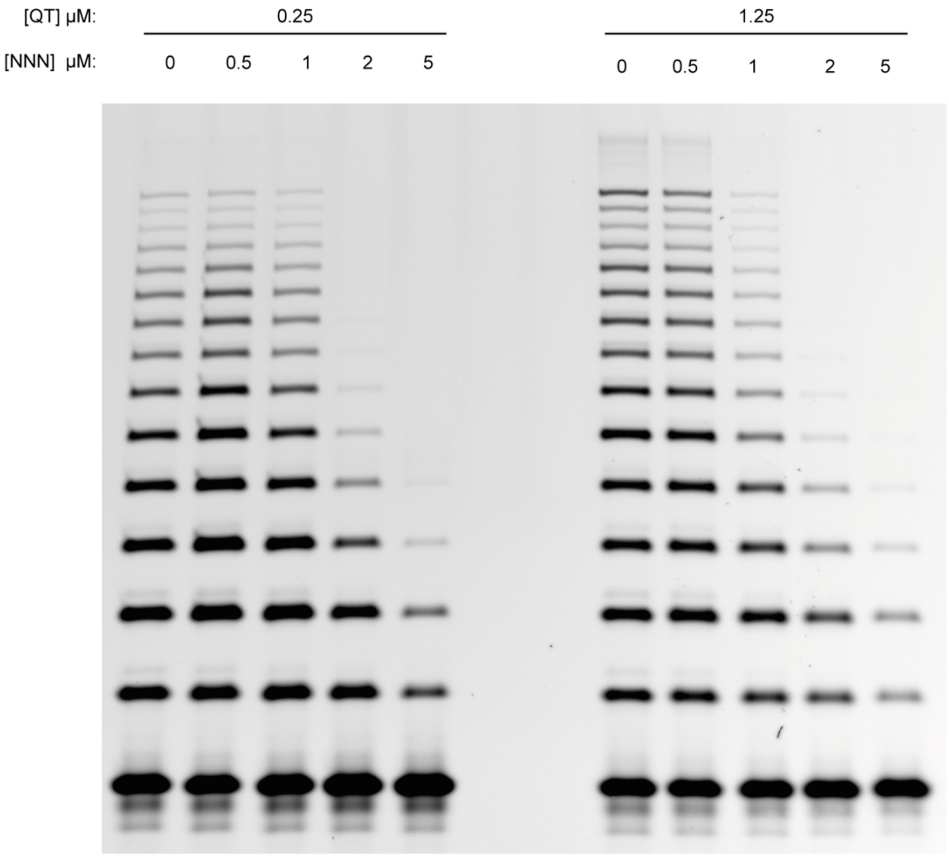


**Fig. S14 Inhibition of ribozyme polymerase activity by high concentration of NNN.**
Synthesis of 42 nucleotide CGU repeat sequence by QT45 at varying concentration of QT45 and NNN (equimolar mix of all possible triphosphorylated trinucleotides). Reaction conditions: 0.25 µM primer F10, 0.25 µM template tP1014CGU, 0.25 µM/1.25 µM ribozyme QT45, 5 µM pppCGU triplet, varying amount of each triplet in pppNNN (shown on gel), 50 mM $MgCl_2$, 50 mM CHES-KOH, pH 9, 3 days at -7 °C frozen.

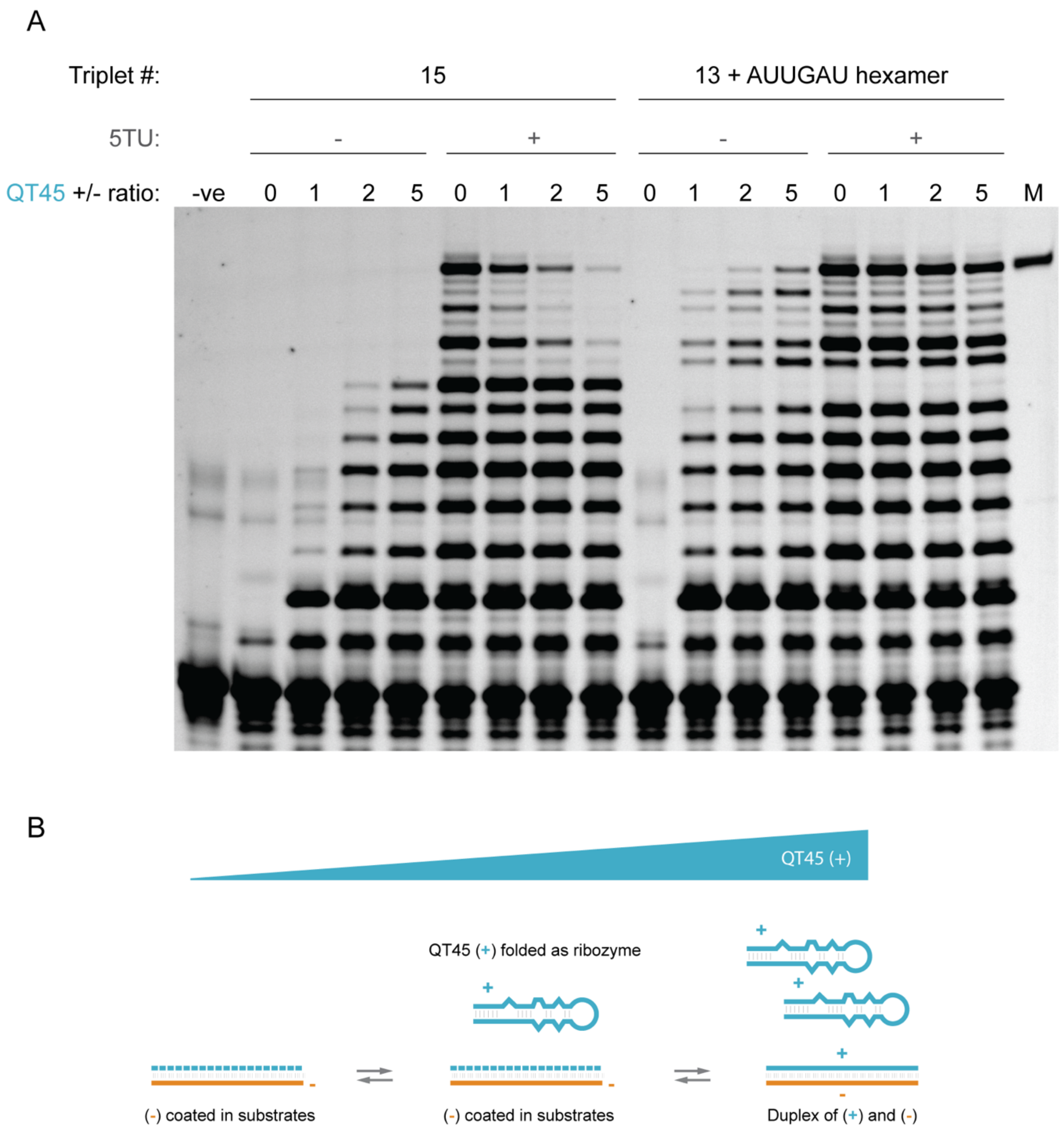


**Fig. S15: Strand inhibition in QT45 (+) strand synthesis.**
(**A**) QT45-catalyzed synthesis of itself (QT45(+)) with increasing ratio of (+) over (-) strand, compared with the same reaction supplemented with 5TU as an additional catalyst. Full-length is indicated by the synthetic marker on the rightmost lane. Reactions were performed using a mix of triplet substrates with or without the aid of one pre-formed hexamer. Reaction conditions: Reaction conditions: 8 nM primer, 8 nM template, 0-40 nM QT45, 0/8 nM 5TU+t1.5, 200 nM each triplet, 0/200 nM pppAUUGAU hexamer, 0.01% Tween 20, 0.4 mM $MgCl_2$, 0.6 mM KCl, 1 mM CHES-KOH, pH 9, acid-heat-cycled once, incubated for 27 days at -7 °C frozen. (**B**) Diagram of the effect of increasing the QT45 (+) strand concentration relative to (-). As the (+) strand increases, more (-) strands are hybridized to their reverse complement, making the template inaccessible by the ribozyme for polymerization.

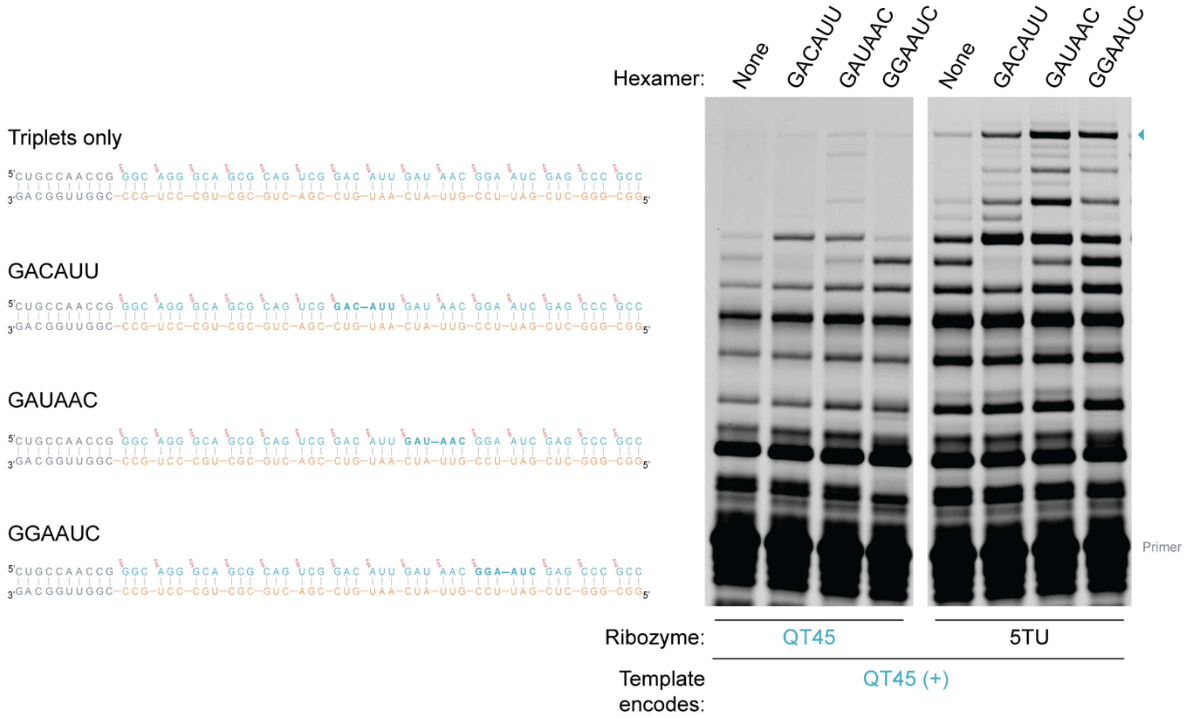


**Fig. S16: Scan of individual hexameric substrates for (+) strand synthesis.**
QT45-catalyzed synthesis of itself (QT45 (+)) with defined triplets and a hexamer compared with the same reaction supplemented with 5TU as an additional catalyst. Full-length is indicated by the triangle in teal. Reactions were performed using a mix of triplet substrates supplemented with a single pre-formed hexamer at various positions. Reaction conditions: 20 nM primer BCy3P10, 8 nM template, 16 nM QT45, 100 nM each triplet and hexamer, 0.01% Tween 20, 0.4 mM $MgCl_2$, 0.6 mM KCl, 1 mM CHES-KOH, pH 9, acid-heat-cycled once, incubated for 28 days at -7 °C frozen.

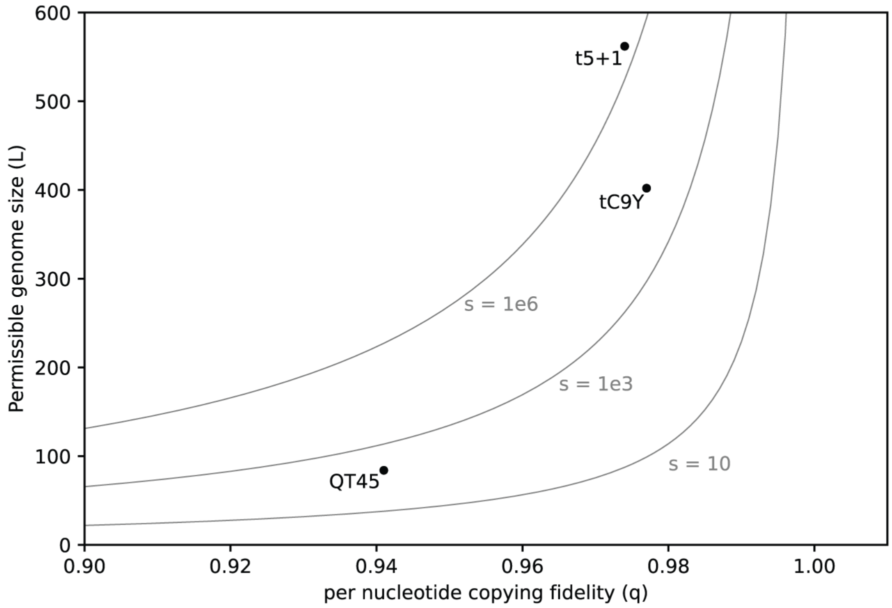


**Fig. S17 Eigen error threshold.**

(**A**) Maximum permissible maximum genome size in nucleotides (*L)* in relation to the fidelity of replication. *L* was calculated using the Eigen error threshold equation: $L = -log(s)/log(q)$ (*6*, *9*). With (*s*) being the selective superiority of the master sequence, which corresponds to the replication rate of the master sequence over the average replication rate of its mutants, and (*q*) being the per nucleotide average replication accuracy. Per nucleotide average fidelity was used for t5+1 and QT45, rather than per trinucleotide, in order to compare more easily with the existing literature. Three different values of selective superiority (*s*) are shown as representative examples, as the selective superiority in a self-replication context is unknown. QT45, tC9Y and t5+1 ribozyme are displayed based on their genome length. This includes the length of both (+) and (-) strand, excluding the first triplet in each strand's synthesis i.e. 84 for QT45, 402 for tC9Y and 568 for t5+1. The fidelity values used are 0.941 for QT45, this manuscript, 0.977 for tC9Y, measured in (*65*), 0.974 for t5+1, measured in (*13*). QT45 would remain below the error threshold for a selective superiority higher than 166, tC9Y would need a selective superiority higher than $1.1 \times 10^4$, and t5+1 would need a selective superiority higher than $2.6 \times 10^6$ to persist.

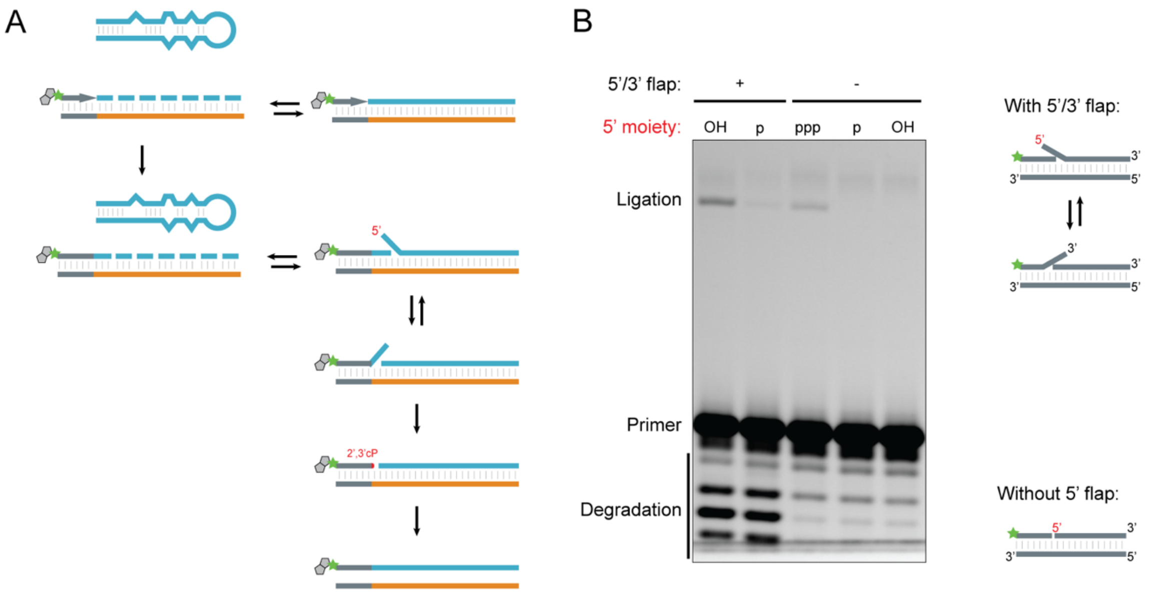


**Fig. S18: Recombination between partial self-synthesis products and the ribozyme.**
(**A**) Model for the recombination between partial self-synthesis products and the ribozyme used for self-synthesis. After incorporation of 1+ triplets the ribozyme can hybridize to the template, leading to a 5′-end flap. This can interchange with the partially extended product, leading to a 3′-end flap. Degradation of this flap leads to the formation of a 2′3′-cyclic phosphate, which can then lead to the templated ligation to the ribozyme used for synthesis that is hybridized to the template (this type of degradation followed by recombination was previously observed in (*46*)). 5′-end phosphorylation can reduce this side reaction, as now two separate degradation events need to occur for ligation to occur. (**B**) Model system for the study of recombination. The presence of a 5′/3′ flap leads to a ligation product that migrates like the product of nonenzymatic ligation to a 5′ triphosphorylated control oligo (a phenomenon previously described in (*66*)). Presence of a monophosphate at the 5′ reduces the extent of recombination. In the absence of the 5′/3′ flap no ligation is observed. Reaction conditions: 0.5 µM primer F10, 0.5 µM template tempF10Ltest1, 0.5 µM substrate (pppLtest1 or pLtest1 or ccgpLtest1 or pccgpLtest1), 0.01% Tween 20, 50 mM $MgCl_2$, 50 mM CHES-KOH, pH 9, incubated for 16 hours at 37 °C and then 7 days at -7 °C frozen.

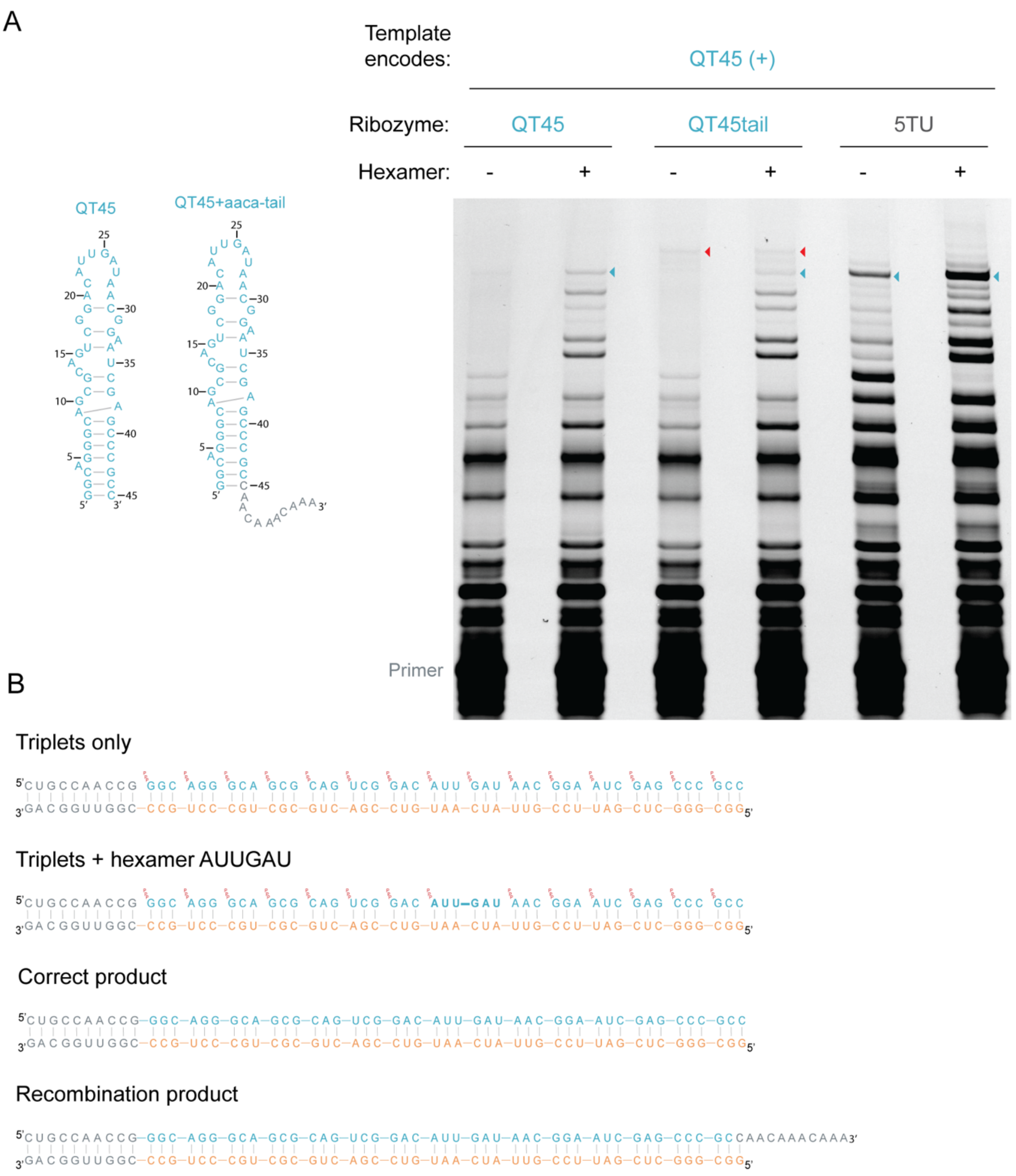


**Fig. S19: QT-ribozyme-catalyzed synthesis of itself.**
(**A**) Synthesis of QT45(+) by either QT45, QT45 with an added 3′-tail, or 5TU. Diagrams of the QT45 and QT45-tail variants used for this experiment are shown on the left, with nucleotides in grey indicating DNA modification. A DNA modification was introduced in the 3′-tail nucleotides to prevent adapter ligation and distinguish synthetic from recombined products via size mobility and sequencing. Reactions were carried out using a mix of triplet substrates with or without the aid of one pre-formed hexamer. The addition of a 3′-tail to QT45 allows us to distinguish the synthetic product from products of recombination with the ribozyme. Full-length is indicated by a teal triangle, incorrect recombination product that is larger and hence migrates with slower mobility is indicated by a red triangle. All sequencing reactions of QT45 (+) strand synthesis described in Fig. 4 were run with these variants to ensure discrimination. Bottom diagrams indicate the reactions without and with the hexamer substrate, the correct synthesis product, as well as the potential recombination product. Reaction conditions: 20 nM primer BCy3P10, 8 nM template, 16 nM QT45, 100 nM each triplet and hexamer, 0.01% Tween 20,

0.4 mM $MgCl_2$, 0.6 mM KCl, 1 mM CHES-KOH, pH 9, acid-heat-cycled once, incubated for 30 days at -7 °C frozen.

**Table S1.** ***De novo*** **selection conditions.**

*N20 from R5_1L and R5_2L were mixed and carried forward as 'N20'. N30 and N40 from R5_2L were mixed and carried forwards as "Mix". Error prone PCR was carried out on all libraries before round 6. ** Libraries were monomeric from round 8 onwards. "Mix" library from previous round was gel purified and its two, originally, N30 and N40 libraries were separated. These were respectively mixed with the N30 and N40 libraries from in the input to round 8. Reaction conditions: 50 nM library/primer/template in round 1, 20 nM in later rounds.

| Round number | DNA used (pmoles) | RNA used (pmoles) | Substrate | Template/primer | Time | % ligated |
|---|---|---|---|---|---|---|
| R1 | ~10 | 1000 | Single ligation | tempF6F10 BTCy3P10 | 73h | 0.1 |
| R2 | ~10 | 100 | Single ligation | tempF6F10 BTCy3P10 | 15.5h | 0.02 |
| R3 | ~6 | 100 | Single ligation | tempF6F10 BTCy3P10 | 13h | 0.03 |
| R4_1L | na | 20 | Single ligation | tempF6F10 BTCy3P10 | 15h | 0.05 |
| R4_2L | na | 20 | 1 triplet (GCG) | temp6FnewP10GCG BCy3newP10 | 15h | na |
| R5_1L | na | 20 | Single ligation | tempF6F10 BTCy3P10 | 16h | 0.94% (N20), 3.26% (N30), 2.25% (N40) |
| R5_2L | na | 20 | 1 triplet (GCG) | temp6FnewP10GCG BCy3newP10 | 16h | 2.47% (N20), 0.15% (N30), 0.28% (N40) |
| R6* | na | 20 | 1 triplet (GCG) | temp6FnewP10GCG BCy3newP10 | 15h | 0.03% (N20), 0.02% (N30), 0.04% (N40), 0.27% (Mix) |
| R7 | na | 20 | 1 triplet (GCG) | temp6FnewP10GCG BCy3newP10 | 16h | 0.2% (N20), 0.03% (N30), 0.27% (N40), 0.54% (Mix) |
| R8** | na | 20 | 3 triplets (GCG) | temp6FP10gaGCG3 BCy3P10GA | 16h | na |
| R9 | na | 20 | 3 triplets (3ACC) | temp6FP10gaACC3 BCy3P10GA | 14h | na |
| R10 | na | 20 | 3 triplets (3ACC) | temp6FP10gaACC3 BCy3P10GA | 17h | na |
| R11 | na | 20 | 3 triplets (3ACC) | temp6FP10gaACC3 BCy3P10GA | 2h | na |

**Table S2. Reselection conditions**

| Round number | RNA used (pmoles) | Substrate | Template/primer | Time |
|---|---|---|---|---|
| R12 | 20000 | 3 triplets (GCG) | temp6FP10gaGCG3 BCy3P10ga | 20 h |
| R13 | 200 | 3 triplets (ACC) | t6FP103ACC BCy3P10 | 16 h |
| R14 | 50 | 3 triplets (UGC) | t6FP103UGC BCy3P10 | 16 h |
| R15 | 20 | 12 triplets (CUA) | t6FP1012CUA BCy3P10 | 41 h |
| R16 | 20 | 12 triplets (CUA) | t6FP1012CUA BCy3P10 | 42 h |
| R17 | 10 | 6 triplets (CUU) | t6FnewP10CUU6 BCy3newP10 | 22 h |
| R18 | 20 | 6 triplets (AUA) | t6FnewP10AUA6 BCy3newP10 | 46 h |

**Table S3. Oligonucleotide sequences**

Oligonucleotide sequences are collated below. RNA is colored in orange, DNA is in black. Any modification is annotated with the supplier's specific code. 'GP' describes in house PAGE purified oligonucleotides, 'RNE' describes QIAGEN RNEasy purification, if not stated the sequences were used as supplied. 'IVT' describes T7 *in vitro* transcribed RNA.

| Application | Name | Source, purification | Sequence (5′-3′) | Notes |
|---|---|---|---|---|
| General fill-in/PCR | 5T7 | Sigma | GATCGATCTCGCCCGCGAAATTAATACGACTCACTATA | |
| | HDVrt | Sigma | CTTCTCCCTTAGCCTACCGAAGTAGCCCAGGTCGGACCGCGAGGAGGTGGAGATGCCATGCCGACCC | |
| Homodimer selection libraries and library generation oligos | pTLT | IDT | /5Phos/GGACAGTCAGGCAGT | |
| | fGG17 | IDT | /5Phos/CAAAACAAACAAACAGG | |
| | ULTc2dN40 | IDT, GP | /5Phos/TTGTTTGTTGGACAGTCAGGCAG/ideoxyU/NNNNNNNNNNNNNNNNNNNNNNNNNNNNNNNNNNNNNNNNCCTGTTTGTTTGTTTTG | |
| | ULTc2dN30 | IDT, GP | /5Phos/TTGTTTGTTGGACAGTCAGGCAG/ideoxyU/NNNNNNNNNNNNNNNNNNNNNNNNNNNNNNCCTGTTTGTTTGTTTTG | |
| | ULTc2dN20 | IDT, GP | /5Phos/TTGTTTGTTGGACAGTCAGGCAG/ideoxyU/NNNNNNNNNNNNNNNNNNNNCCTGTTTGTTTGTTTTG | |
| | 5T76FfGG | IDT | GATCGATCTCGCCCGCGAAATTAATACGACTCACTATAGGTCCAAACAAACAACAAAACAAACAAACAGG | |
| | bio5T76FfGG | IDT | /5Biosg/GATCGATCTCGCCCGCGAAATTAATACGACTCACTATAGGTCCAAACAAACAACAAAACAAACAAACAGG | |
| | 5T76FfGGLaaca15cagg | IDT | GATCGATCTCGCCCGCGAAATTAATACGACTCACTATAGGTCCAAACAAACAAACAAACAAACAAACAAAC | Longer version of 5T76FfGG for 15AACA linker |

| | | | | |
|---|---|---|---|---|
| | | | AAACAAACAAACAAACAAACAACAAAACAAACAGG | |
| Reselection libraries | 1-30-sp24 | | CAGGCAGTAAGCAGTGCGTTTTTTACGTTAATTGTTCACCTGTTTGTTTGTTTTGTTGTTTGTTTGGACC | Underlined nt were spiked at 24% |
| | 2-30-sp24 | | GGACAGTCAGGCAGTTACTCGTTAGGTACTTCTTAATTTTTCGCCCCTGTTTGTTTGTTTTGTTGTTTGTTTGGACC | Underlined nt were spiked at 24% |
| | 1-40-sp24 | | CAGGCAGTCTCTCGAGTCCGTTATCTATTTCCGCCTGCGCTGAGATGCCTGTTTGTTTGTTTTGTTGTTTGTTTGGACC | Underlined nt were spiked at 24% |
| Primers and templates used in selection | BTCy3P10 | IDT, GP | /5BiotinTEG//iCy3/CUGCCAACCG | |
| | BCy3P10 | IDT, GP | /5Biosg//iCy3/CUGCCAACCG | |
| | BCy3P10GA | IDT, GP | /5Biosg//iCy3/CUGCCAACCGGA | |
| | BCy3newP10 | IDT, GP | /5BiosG//iCy3/CGCACUCAGG | |
| | temp6FF10 | IDT, GP | UGGACCCGGUUGGCAG/3SpC3/ | |
| | temp6FnewP10GCG | IDT, GP | UGGACCCGCCCUGAGUGCG/3SpC3/ | |
| | t6FP10gaGCG3 | IDT, GP | UGGACCCGCCGCCGCUCCGGUUGGCAG/3SpC3/ | |
| | temp6FP10gaACC3 | IDT, GP | UGGACCGGUGGUGGUUCCGGUUGGCAG/3SpC3/ | |
| | temp6FP10ACC3 | IDT, GP | UGGACCGGUGGUGGUCGGUUGGCAG/3SpC3/ | |
| | temp6FP103ugc | IDT, GP | UGGACCgcagcagcaCGGUUGGCAG/3SpC3/ | |
| | temp6FP1012cua | IDT, GP | UGGACCuaguaguaguaguaguaguaguaguaguaguagCGGUUGGCAG | |
| | temp6FnewP10CUU6 | IDT, GP | UGGACCaagaagaagaagaagaagCCUGAGUGCG/3SpC3/ | |
| | temp6FnewP10AUA6 | IDT, GP | UGGACCuauuauuauuauuauuauCCUGAGUGCG/3SpC3/ | |
| Recovery primers | HDVrec | IDT | GATGCCATGCCGACCC | Used for recovery of rounds 12-18 alongside round-specific primers described below |

| | | | | |
|---|---|---|---|---|
| | pP106Frec | IDT | /5Phos/CTGCCAACCGGG TCCA | Used for recovery of rounds R1, R2, R3, R4_1L alongside pTLT |
| | pnewP10gcFrec | IDT | /5Phos/CGCACTCAGGGC | Used for recovery of rounds R4_2L, R5_1L, R5_2L, R6, R7 alongside pTLT |
| | pP10GAgcgFrec | IDT | /5Phos/TGCCAACCGGAgc | Used for recovery of rounds R8, R12 |
| | pP10GAaccFrec | IDT | /5Phos/CTGCCAACCGGA ac | Used for recovery of rounds R9, R10, R11 |
| | P10accFrec | IDT | CTGCCAACCGacc | Used for recovery of R13 |
| | P10ugcFrec | IDT | CTGCCAACCGtgc | Used for recovery of R14 |
| | forceGG | Sigma | AACAAACAACAAAACAAA CAAACAGG | Used for recovery in R15, R16, R17, R18 |
| Adapters used in recovery | AdeHDVlig | | Ap-pGGGTCGGCATGGCATC/ 3SpC3/ | Prepared via adenylation as in method 1.4, starting from HDVlig |
| | HDVlig | IDT, GP | /5Phos/GGGTCGGCATGG CATC/3SpC3/ | |
| Regiospecificity assay | FITCreg3P | | FITC-CAAUACAACCG-3p(-2p) | Marker for cleaved reaction |
| | FITCreg3 | | FITC-CAAUACAACCG | Generated via T4 PNK dephosphorylation of FITCreg3P |
| | t6Freg3GCG2CUG | IDT | UGGACCCAGCGCCGCC GGUUGUAUUG | Template for regiospecificity assay. |
| | ohCUG | ChemGenes | ohCUG | Monophosphorylated chemically synthesized triplet |
| Clones from de novo selection | 8_R11_20 (1-30) | IVT, GP | GGUCCAAACAAACAACA AAACAAACAAA**CAGGUG AACAAUUAACGUAAAAA ACGCACUGCUUACUGC CUG** | Underlined nucleotides hybridize to the template. Bold nucleotides indicate residues likely to fold into ribozyme. |

| | | | | |
|---|---|---|---|---|
| | 8_R11_20_R | IDT | CAGGCAGTAAGCAGTGCGTTTTTTACGTTAATTGTTCACCTGTTTGTTTGTTTTGTTGTTTGTTTGGACC | Used together with 5T76FfGG to generate template for transcription of 8_R11_r0 |
| | 0_R8_30 (2-30) | IVT, GP | GGUCCAAACAAACAACAAAACAAACAAA**CAGGGGCGAAAAAUUAAGAAGUACCUAACGAGUAACUGCCUGACUGUCC** | Underlined nucleotides hybridize to the template. Bold nucleotides indicate residues likely to fold into ribozyme. |
| | 0_R8_30_R | IDT | GGACAGTCAGGCAGTTACTCGTTAGGTACTTCTTAATTTTTCGCCCCTGTTTGTTTGTTTTGTTGTTTGTTTGGACC | Used together with 5T76FfGG to generate template for transcription of 1_R11_40 |
| | 1_R11_40 (1-40) | IVT, GP | GGUCCAAACAAACAACAAAACAAACAAA**CAGGCAUCUCAGCGCAGGCGGAAAUAGAUAACGGACUCGAGAGACUGCCUG** | Underlined nucleotides hybridise to the template. Bold nucleotides indicate residues likely to fold into ribozyme. |
| | 1_R11_40_R | IDT | CAGGCAGTCTCTCGAGTCCGTTATCTATTTCCGCCTGCGCTGAGATGCCTGTTTGTTTGTTTTGTTGTTTGTTTGGACC | Used together with 5T76FfGG to generate template for transcription of 1_R11_40 |
| Ribozymes used in this study | 5TU | IVT,GP | GGAUCUUCUCGAUCUAACAAAAAAGACAAAUCUGCCACAAAGCUUGAGAGCAUCUUCGGAUGCAGAGGCGGCAGCCUUCGGUGGCGCGAUAGCGCCAACGUUCUCAACUAUGACACGCAAAACGCGUGCUCCGUUGAAUGGAGUUUAUCAUG | |
| | 5T7_5TUF | Sigma | GATCGATCTCGCCCGCGAAATTAATACGACTCACTATAGGATCTTCTCGATCTAACAAAAAAGACAAATCTGCCACAAAGCTTGAGAGCATCTTCGGATG | Used with 5TUR for fill-in and generate a template for 5TU |

| | | | | |
|---|---|---|---|---|
| | 5TUR | Sigma | CATGATAAACTCCATTCA ACGGAGCACGCGTTTTG CGTGTCATAGTTGAGAA CGTTGGCGCTATCGCGC CACCGAAGGCTGCCGCC TCTGCATCCGAAGATGC TCTCAAGCTTTGTGGCA | |
| | T1.5 | IVT, GP | GACCAAUCUGCCCUCAG AGCCCGAGAACAUCUUC GGAUGCAGAGGAGGCA GGCUUCGGUGGCGCGA UAGCGCCAACGUCCUCA ACCUCCAAUGCAUCCCA CCACAUGAUGAGCCUGA AGAGCCUUGGUUUUUUU G | |
| | 5T7_t1.5F | Sigma | GATCGATCTCGCCCGCG AAATTAATACGACTCACT ATAGACCAATCTGCCCTC AGAGCCCGAGAACATCT TCGGATGCAGAGGAG | Used with t1.5R to fill-in and generate a template for t1.5 |
| | t1.5R | Sigma | CAAAAAAACCAAGGCTC TTCAGGCTCATCATGTG GTGGGATGCATTGGAGG TTGAGGACGTTGGCGCT ATCGCGCCACCGAAGCC TGCCTCCTCTGCATCCG | |
| | QT51 | IDT, GP | ACAGGCAUCUCAGCGCA GUCGGACAUUGAUAAUG GAAUCGAGAGAGUCUGU | |
| | QT45 | IDT, GP | GGCAGGGCAGCGCAGU CGGACAUUGAUAACGGA AUCGAGCCCGCC | |
| | pQT45 | IDT,GP | /5Phos/GGCAGGGCAGCG CAGUCGGACAUUGAUAA CGGAAUCGAGCCCGCC | 5′-end monophosphate used to reduce recombination to partial synthesis products (see Fig. S17) |
| | QT40 | IDT, GP | GGGGCAGCGCAGUCGG ACAUUGAUAACGGAAUC GAGCCCC | |
| | QT35 | IDT, GP | GGCAGCGCAGUCGGAA UUGAUAAUGGAAUCGAG CC | |
| | QT45aaca | | GGCAGGGCAGCGCAGU CGGACAUUGAUAACGGA AUCGAGCCCGCC AACAAACAAA | 3′-end DNA tail use to distinguish synthetic products from QT ribozyme used for |

| | | | | |
|---|---|---|---|---|
| | | | | synthesis. Used in Fig. S18 and in sequencing of (+) strand synthesis. |
| | QT39 | IVT, GP | GGGGCAGCGCAGUCGGACAUUGAUAACGGAAUCGAGCCC | Used in Fig. S8 |
| | 0_51_r7trim | IDT, GP | GGUCCAAACAAACAACAAAACAAACAAA**GGCCAGCGCAGTCGGACATTGATAATGGAATCGAGGCC** | Underlined nucleotides hybridize to the template. Bold nucleotides indicate residues likely to fold into ribozyme. Used in fig. S7 |
| | 0_qt51_r7_trim_R | IDT | GGCCTCGATTCCATTATCAATGTCCGACTGCGCTGGCCTTTGTTTGTTTTGTTGTTTGTT | Used with 5T76FnocaGG (below) to prepare the template for transcription of 0_51_r7trim |
| | 5T76FnocaGG | IDT | GATCGATCTCGCCCGCGAAATTAATACGACTCACTATAGGTCCAAACAAACAACAAAACAAACAAAGG | |
| | QT51_6F5L | IVT, GP | GGUCCAAACAACAAAACAAACAAA**ACAGGCAUCUCAGCGCAGUCGGACAUUGAUAAUGGAAUCGAGAGAGUCUGU** | Underlined nucleotides hybridize to the template. Bold nucleotides indicate residues likely to fold into ribozyme. Used in fig. S6 |
| | QT51_6F5L_F | | GATCGATCTCGCCCGCGAAATTAATACGACTCACTATAGGTCCAACAAAACAAACAAACAGGCATCTCAGC | Used with QT51_6F5L_R for fill-in to generate the template for transcription of QT51_6F5L |
| | QT51_6F5L_R | | ACAGACTCTCTCGATTCCATTATCAATGTCCGACTGCGCTGAGATGCCTGTTTGTTTGTTTTG | |
| Primers used for RNA-catalysed | F10 | IDT, GP | /56-FAM/CUGCCAACCG | |

| | | | | |
|---|---|---|---|---|
| RNA synthesis | | | | |
| Templates used for RNA-catalysed RNA synthesis | tP10CGU20 | IVT, GP | GGACGACGACGACGACGACGACGACGACGACGACGACGACGACGACGACGACGACGACGACGCGGUUGGCAG | Primer binding site underlined. |
| | tP10_20cgu(tx) | IDT | CTGCCAACCGCGTCGTCGTCGTCGTCGTCGTCGTCGTCGTCGTCGTCGTCGTCGTCGTCGTCGTGGTATAGTGAGTCGTATTAATTTCGC | Used with 5T7 to prepare the template for transcription of tP10CGU20. |
| | P10CGU20comp | IVT, RNE | GGCUGCCAACCGCGUCGUCGUCGUCGUCGUCGUCGUCGUCGUCGUCGUCGUCGUCGUCGUCGUCGUCGU | Used as “competing oligo” to improve resolution of tP10CGU20 copying reaction on a denaturing PAGE. |
| | P10CGU20comp(tx) | IDT | ACGACGACGACGACGACGACGACGACGACGACGACGACGACGACGACGACGACGACGACGCGGTTGGCAGCCTATAGTGAGTCGTATTAATTTCGCGGGC | Used with 5T7 to prepare the template for transcription of P10CGU20comp. |
| | tP10CGU14 | IDT, GP | ACGACGACGACGACGACGACGACGACGACGACGACGACGACGCGGUUGGCAG | Primer binding site underlined. |
| | P10CGU14comp | IVT, RNE | GGCUGCCAACCGCGUCGUCGUCGUCGUCGUCGUCGUCGUCGUCGUCGUCGU | Used as “competing oligo” to improve resolution of tP10CGU14 copying reaction on a denaturing PAGE. |
| | P10CGU14comp(tx) | IDT | ACGACGACGACGACGACGACGACGACGACGACGACGACGCGGTTGGCAGCCTATAGTGAGTCGTATTAATTTCGCGGGCGAGATCGA | Used with 5T7 to prepare the template for transcription of P10CGU14comp |
| | t6FP10mix | IDT, GP | UGGACCUAUGCGUUCGAAGGUCGCCGGUUGGCAG/3SpC3/ | |
| | rc_of_t6FP10mix | IDT, GP | CUGCCAACCGGCGACCUUCGAACGCAUAGGUCCA | Used as “competing oligo” |

| | | | | |
|---|---|---|---|---|
| | | | | to improve resolution of t6FP10mix copying reaction on a denaturing PAGE. |
| Mini hammerhead synthesis, activity, sequencing | tP10HHz | IDT, GP | GCGCCUCAUCAGUCGAG CCGGUUGGCAG | Template used for mini hammerhead synthesis in Fig. 3 |
| | Fsubuhl | IDT, GP | /56-FAM/GCGCCGAAACACC GUGUCUCGAGC | Substrate for hammerhead cleavage |
| | DNAtP10_Euhlp1 | IDT | GCGCCTCATCAGTCGAG CCGGTTGGCAG | Template used for TGK synthesis of hammerhead marker |
| | A647BP10 | IDT | /5Alex647N//iBiodT/CUGC CAACCG | Primer used for synthesis of mini hammerhead used for cleavage assay |
| | bioCy3KyleP10 | IDT, GP | /5BiosG//iCy3/GGATTCAC TGCGATAGAGTCCUGCC AACCG | Primer used for synthesis of mini hammerhead used for sequencing |
| Templates used for (+) and (-) strand synthesis | t4psP10QT45 | IDT, GP | GGCGGGCUCGAUUCCG UUAUCAAUGUCCGACUG CGCUGCCCUGCCCGGU UGGCAG | Template for (+) strand synthesis in Fig. 4 |
| | t4msP10QT45 | IDT, GP | GCAGGGCAGCGCAGUC GGACAUUGAUAACGGAA UCGAGCCCGCCCCGGU UGGCAG | Template for (-) strand synthesis in Fig. 4 |
| Hexamer synthesis | pppGACAUU | IVT, GP | pppGACAUU | |
| | GACAUU_tx | Sigma | AATGTCTATAGTGAGTCG TATTAATTTCGCGGGCGA GATCGATC | |
| | pppGAUAAC | IVT, GP | pppGAUAAC | |
| | GAUAAC_tx | Sigma | GTTATCTATAGTGAGTCG TATTAATTTCGCGGGCGA GATCGATC | |
| | pppGGAAUC | IVT, GP | pppGGAAUC | |
| | GGAAUC_tx | Sigma | GATTCCTATAGTGAGTCG TATTAATTTCGCGGGCGA GATCGATC | |
| | pppAUUGAU | Chemgenes, GP | pppAUUGAU | |

| | | | | |
|---|---|---|---|---|
| Analysis of recombination | pppLtest1 | IVT, GP | pppGCGAAGCGUGU | Used in fig. S17 as triphosphorylated |
| | (tx)Ltest1 | IDT | ACACGCTTCGCTATAGTGAGTCGTATTAATTTCGCGGGCGAGATCGATC | Fill-in with 5T7 to generate template for pppLtest1 |
| | Ltest | IDT, GP | GCGAAGCGUGU | Used in fig. S17 |
| | pLtest | IDT, GP | /5Phos/GCGAAGCGUGU | Used in fig. S17 |
| | AppLtest | GP | App-GCGAAGCGUGU | Adenylated pLtest with method 1.4 |
| | ccgpLtest1 | IDT, GP | CCGGCGAAGCGUGU | Used in fig. S17 |
| | pccgpLtest1 | IDT, GP | /5Phos/CCGGCGAAGCGUGU | Used in fig. S17 |
| Deoxynucleotide substitution scanning | F10_d-1 | Sigma, GP | /56-FAM/CUGCCAACC[dG] | |
| | F10_d-2 | Sigma, GP | /56-FAM/CUGCCAAC[dC]G | |
| | F10_d-3 | Sigma, GP | /56-FAM/CUGCCAA[dC]CG | |
| | F10_d-4 | Sigma, GP | /56-FAM/CUGCCA[dA]CCG | |
| | F10_d-5 | Sigma, GP | /56-FAM/CUGCC[dA]ACCG | |
| | tF10Ltest_d-1 | Sigma, GP | ACACGCUUCGC[dC]GGUUGGCAG | |
| | tF10Ltest_d-2 | Sigma, GP | ACACGCUUCGCC[dG]GUUGGCAG | |
| | tF10Ltest_d-3 | Sigma, GP | ACACGCUUCGCCG[dG]UUGGCAG | |
| | tF10Ltest_d-5 | Sigma, GP | ACACGCUUCGCCGG[dU]UGGCAG | |
| | tF10Ltest_d+1 | Sigma, GP | ACACGCUUCG[dC]CGGUUGGCAG | |
| | tF10Ltest_d+2 | Sigma, GP | ACACGCUUC[dG]CCGGUUGGCAG | |
| | tF10Ltest_d+3 | Sigma, GP | ACACGCUU[dC]GCCGGUUGGCAG | |
| | tF10Ltest_d+4 | Sigma, GP | ACACGCU[dU]CGCCGGUUGGCAG | |
| | tF10Ltest_d+5 | Sigma, GP | ACACGC[dU]UCGCCGGUUGGCAG | |
| | tF10Ltest_GCGtGCA | Sigma, GP | ACACGCUUUGCCGGUUGGCAG | |
| | DSsub | IVT, GP | pppGCGAAGCGUGU | |
| | tDSsub | Sigma | ACACGCTTCGCTATAGTGAGTCGTATTAATTTCGC | Fill-in with 5T7 to generate template for DSsub |

| | | | | |
|---|---|---|---|---|
| | DSsub_GCGtGCA | IVT, GP | pppGCAAAGCGUGU | |
| | tDSsub_GCGtGCA | Sigma | ACACGCTTTGCTATAGTG AGTCGTATTAATTTCGC | Fill-in with 5T7 to generate template for DSub_GCGtGCA |
| | DSsub_GCGtGCA_d +1 | Chemge nes, GP | ppp[dG]CAAAGCGUGU | |
| | DSsub_GCGtGCA_d +2 | Chemge nes, GP | pppG[dC]AAAGCGUGU | |
| | DSsub_GCGtGCA_d +3 | Chemge nes, GP | pppGC[dA]AAGCGUGU | |
| | DSsub_GCGtGCA_d +4 | | pppGCA[dA]AGCGUGU | Prepared via splinted ligation of pppGCA to DSsub_minGCA_ d+4 |
| | DSsub_minGCA_d+ 4 | IDT, GP | [dA]AGCGUGU | |
| | DSsub_GCGtGCA_d +5 | | pppGCAA[dA]GCGUGU | Prepared via splinted ligation of pppGCA to DSsub_minGCA_ d+5 |
| | DSsub_minGCA_d+ 5 | IDT, GP | A[dA]GCGUGU | |
| | tTripletLigation_3spc 3 | IDT, GP | AACAAACAAACAACACG CTTTGC/3SpC3 | Splint used for the generation of DSsub_minGCA_ d+4 and DSsub_minGCA_ d+5 |
| Fitness landscape | P71forceGG_2024 | IDT | CAAGCAGAAGACGGCAT ACGAGATGTGACTGGAG TTCAGACGTGTGCTCTTC CGATCTNNNgatactAACA AACAACAAAACAAACAAA CAGG | Fitness landscape sequencing primer |
| | P72forceGG_2024 | IDT | CAAGCAGAAGACGGCAT ACGAGATGTGACTGGAG TTCAGACGTGTGCTCTTC CGATCTNNNatcttgAACAA ACAACAAAACAAACAAAC AGG | Fitness landscape sequencing primer |
| | P51HDVba_2021 | IDT | AATGATACGGCGACCAC CGAGATCTACACTCTTTC CCTACACGACGCTCTTC CGATCTNNNatcacgGATG CCATGCCGACCC | Fitness landscape sequencing primer |

| | | | | |
|---|---|---|---|---|
| | P52HDVba_2021 | IDT | AATGATACGGCGACCACCGAGATCTACACTCTTTCCCTACACGACGCTCTTCCGATCTNNNcgatgtGATGCCATGCCGACCC | Fitness landscape sequencing primer |
| | P53HDVba_2021 | IDT | AATGATACGGCGACCACCGAGATCTACACTCTTTCCCTACACGACGCTCTTCCGATCTNNNttaggcGATGCCATGCCGACCC | Fitness landscape sequencing primer |
| | P512HDVba_2021 | IDT | AATGATACGGCGACCACCGAGATCTACACTCTTTCCCTACACGACGCTCTTCCGATCTNNNcttgtaGATGCCATGCCGACCC | Fitness landscape sequencing primer |
| | P514HDVba_2021 | IDT | AATGATACGGCGACCACCGAGATCTACACTCTTTCCCTACACGACGCTCTTCCGATCTNNNagttccGATGCCATGCCGACCC | Fitness landscape sequencing primer |
| | P10UGCugFrec | IDT | CTGCCAACCGTGCTG | Fitness landscape construct recovery |
| | forceGG | IDT | AACAAACAACAAAACAAACAAACAGG | Fitness landscape construct recovery |
| | newnewP12auaaFrec | IDT | CGCACGAGTCTCATAA | Fitness landscape construct recovery |
| | BCy3newnewP12 | IDT, GP | /5Biosg//iCy3/CGCACGAGUCUC | Primer used in fitness landscape |
| | BCy3P10 | IDT, GP | /5Biosg//iCy3/CUGCCAACCG | Primer used in fitness landscape |
| | temp6FP10UGC3 | IDT, GP | UGGACCGCAGCAGCACGGUUGGCAG/3SpC3/ | Template used in fitness landscape |
| | temp6FnewnewP12AUA3 | IDT, GP | UGGACCUAUUAUUAUGAGACUCGUGCG/3SpC3/ | Template used in fitness landscape |
| | temp6FP10CUA12 | IDT, GP | UGGACCUAGUAGUAGUAGUAGUAGUAGUAGUAGUAGUAGUAGCGGUUGGCAG | Template used in fitness landscape |
| | QT45MO10 | IDT | /5Phos/GATGCCATGCCGACCCGGCGGGCTCGATTCCGTTATCAATGTCCGAC | Underlined nt were spiked at 12% |

| | | | | |
|---|---|---|---|---|
| | | | TGCGCTGCCCTGCCCTGTTTGTTTGTTTTGTTG | |
| | QT45MO10_iG1_dC45 | IDT | /5Phos/GATGCCATGCCGACCCGCGGGCTCGATTCCGTTATCAATGTCCGACTGCGCTGCCCTGCCCCTGTTTGTTTGTTTTGTTG | Underlined nt were spiked at 12% |
| | QT45MO10_dG1_iC45 | IDT | /5Phos/GATGCCATGCCGACCCGGGCGGGCTCGATTCCGTTATCAATGTCCGACTGCGCTGCCCTGCCTGTTTGTTTGTTTTGTTG | Underlined nt were spiked at 12% |
| | QT45MO10_C21D_ins1G | IDT | /5Phos/GATGCCATGCCGACCCGGCGGGCTCGATTCCGTTATCAATTCCGACTGCGCTGCCCTGCCCCTGTTTGTTTGTTTTGTTG | Underlined nt were spiked at 12% |
| | QT45MO10_C21D_ins45C | IDT | /5Phos/GATGCCATGCCGACCCGGGCGGGCTCGATTCCGTTATCAATTCCGACTGCGCTGCCCTGCCCTGTTTGTTTGTTTTGTTG | Underlined nt were spiked at 12% |
| | QT45MO10_C21D_ins20G | IDT | /5Phos/GATGCCATGCCGACCCGGCGGGCTCGATTCCGTTATCAATTCCCGACTGCGCTGCCCTGCCCTGTTTGTTTGTTTTGTTG | Underlined nt were spiked at 12% |
| | QT45MO10_C21D_ins22G | IDT | /5Phos/GATGCCATGCCGACCCGGCGGGCTCGATTCCGTTATCAACTTCCGACTGCGCTGCCCTGCCCTGTTTGTTTGTTTTGTTG | Underlined nt were spiked at 12% |
| | QT45MO10_U23D_ins45C | IDT | /5Phos/GATGCCATGCCGACCCGGGCGGGCTCGATTCCGTTATCATGTCCGACTGCGCTGCCCTGCCCTGTTTGTTTGTTTTGTTG | Underlined nt were spiked at 12% |
| | QT45MO10_U23D_ins1G | IDT | /5Phos/GATGCCATGCCGACCCGGCGGGCTCGATTCCGTTATCATGTCCGACTGCGCTGCCCTGCCCCTGTTTGTTTGTTTTGTTG | Underlined nt were spiked at 12% |

| | | | | |
|---|---|---|---|---|
| | HDVrest | IDT | /5AmMC6/CTTCTCCCTTAGCCTACCGAAGTAGCCCAGGTCGGACCGCGAGGAGGTGGA | Used to ligate to library oligos and add the HDV sequence |
| | HDVspl | IDT | /5AmMC6/GGGTCGGCATGGCATCTCCACCTCCTCG/3AmMO/ | Used to split the ligation of library oligos and add the HDV sequence |
